# Differential chromatin accessibility and transcriptional dynamics define breast cancer subtypes and their lineages

**DOI:** 10.1101/2023.10.31.565031

**Authors:** Michael D. Iglesia, Reyka G. Jayasinghe, Siqi Chen, Nadezhda V. Terekhanova, John M. Herndon, Erik Storrs, Alla Karpova, Daniel Cui Zhou, Nataly Naser Al Deen, Andrew T. Shinkle, Rita Jui-Hsien Lu, Wagma Caravan, Andrew Houston, Yanyan Zhao, Kazuhito Sato, Preet Lal, Cherease Street, Fernanda Martins Rodrigues, Austin N. Southard-Smith, André Luiz N. Targino da Costa, Houxiang Zhu, Chia-Kuei Mo, Lisa Crowson, Robert S. Fulton, Matthew A. Wyczalkowski, Catrina C. Fronick, Lucinda A. Fulton, Hua Sun, Sherri R. Davies, Elizabeth L. Appelbaum, Sara E. Chasnoff, Madelyn Carmody, Candace Brooks, Ruiyang Liu, Michael C. Wendl, Clara Oh, Diane Bender, Carlos Cruchaga, Oscar Harari, Andrea Bredemeyer, Kory Lavine, Ron Bose, Julie Margenthaler, Jason M. Held, Samuel Achilefu, Foluso Ademuyiwa, Rebecca Aft, Cynthia Ma, Graham A. Colditz, Tao Ju, Stephen T. Oh, James Fitzpatrick, E. Shelley Hwang, Kooresh I. Shoghi, Milan G. Chheda, Deborah J. Veis, Feng Chen, Ryan C. Fields, William E. Gillanders, Li Ding

## Abstract

Breast cancer is a heterogeneous disease, and treatment is guided by biomarker profiles representing distinct molecular subtypes. Breast cancer arises from the breast ductal epithelium, and experimental data suggests breast cancer subtypes have different cells of origin within that lineage. The precise cells of origin for each subtype and the transcriptional networks that characterize these tumor-normal lineages are not established. In this work, we applied bulk, single-cell (sc), and single-nucleus (sn) multi-omic techniques as well as spatial transcriptomics and multiplex imaging on 61 samples from 37 breast cancer patients to show characteristic links in gene expression and chromatin accessibility between breast cancer subtypes and their putative cells of origin. We applied the PAM50 subtyping algorithm in tandem with bulk RNA-seq and snRNA-seq to reliably subtype even low-purity tumor samples and confirm promoter accessibility using snATAC. Trajectory analysis of chromatin accessibility and differentially accessible motifs clearly connected progenitor populations with breast cancer subtypes supporting the cell of origin for basal-like and luminal A and B tumors. Regulatory network analysis of transcription factors underscored the importance of BHLHE40 in luminal breast cancer and luminal mature cells, and KLF5 in basal-like tumors and luminal progenitor cells. Furthermore, we identify key genes defining the basal-like (*PRKCA*, *SOX6*, *RGS6*, *KCNQ3*) and luminal A/B (*FAM155A*, *LRP1B*) lineages, with expression in both precursor and cancer cells and further upregulation in tumors. Exhausted CTLA4-expressing CD8+ T cells were enriched in basal-like breast cancer, suggesting altered means of immune dysfunction among breast cancer subtypes. We used spatial transcriptomics and multiplex imaging to provide spatial detail for key markers of benign and malignant cell types and immune cell colocation. These findings demonstrate analysis of paired transcription and chromatin accessibility at the single cell level is a powerful tool for investigating breast cancer lineage development and highlight transcriptional networks that define basal and luminal breast cancer lineages.

## INTRODUCTION

Breast cancer (BC) is the most common cancer in women, with 2.1 million new cases diagnosed in 2018^1,2^. Treatment is guided by biomarker profiles, specifically the expression of estrogen receptor (ER), progesterone receptor (PR), and human epidermal growth factor receptor 2 (HER2), which approximate the breast cancer molecular subtypes^3^. Breast ductal epithelium, from which BC arises, is divided into two main lineages (**Fig. 1a**). Luminal cells line the interior of the breast duct, and are surrounded by a layer of thin, contractile basal myoepithelial cells. Both luminal and basal cells are derived from a long-lived, bipotent mammary stem cell, and more differentiated unipotent progenitor cells exist within the basal and luminal lineages to renew these compartments in the breast duct^4,5^. Several groups have interrogated normal and breast cancer cell types at the single-cell level, further refining our understanding of the expression profiles associated with these cell lineages^6–19^.

**Figure 1.**
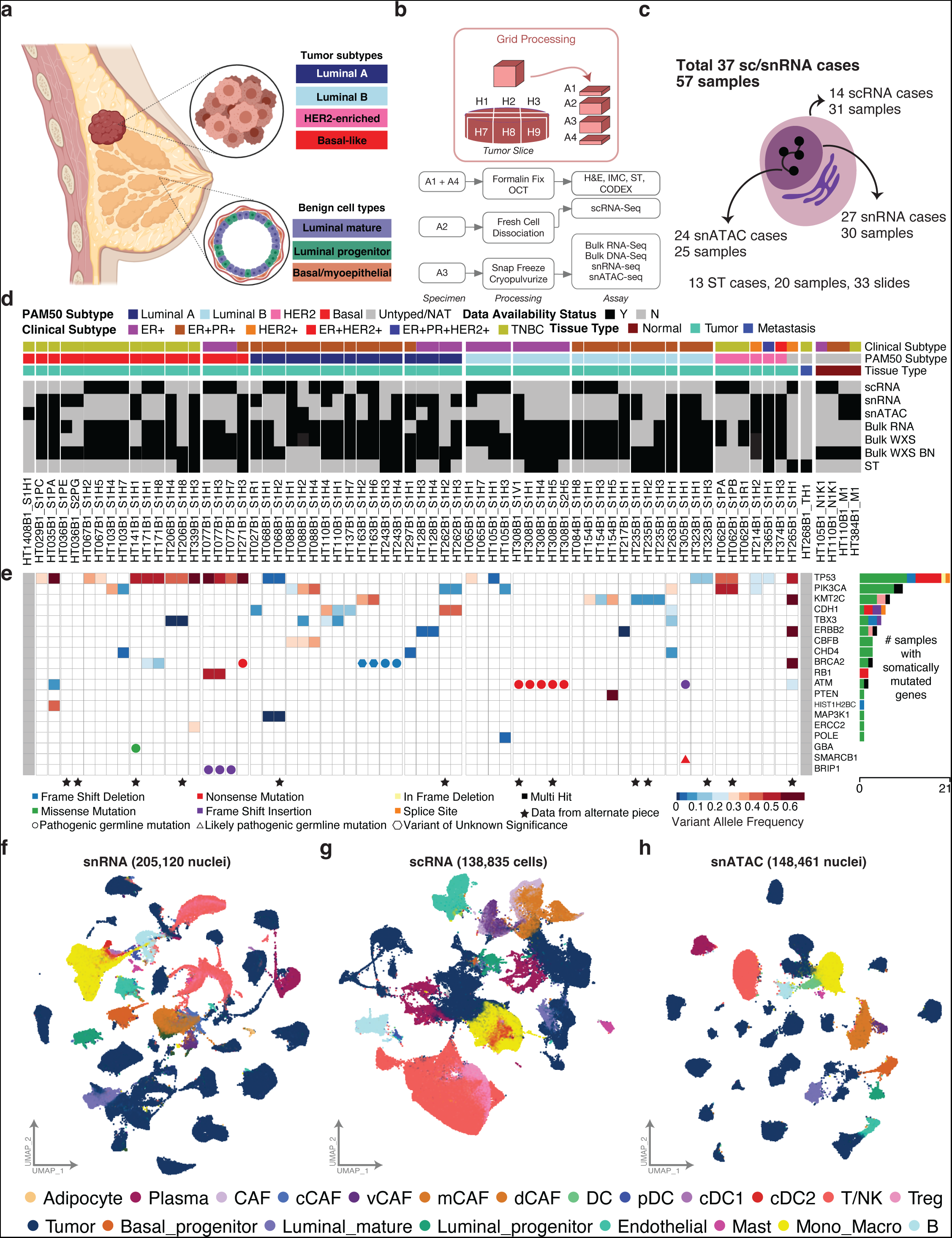
Study Design, Data Collected, and Genomic Alterations. **a)** Summary of benign breast duct cell types and breast cancer subtypes. **b)** Sample grid processing method utilized in the study to perform various assays on each tumor sample systematically. **c)** Summary of data types available for single-cell, single-nucleus and spatial transcriptomics processing. **d)** Data overview of the cohort of 61 samples. The N1K1 and M1 suffix denotes normal adjacent tumor samples. Clinical characteristics and data type availability are shown for each tumor piece. Data types include single-cell RNA-seq (scRNA), single-nuclei RNA-seq (snRNA), single-nuclei ATAC-seq (snATAC), bulk RNA-seq, spatial transcriptomics (ST), and bulk whole exome seq (WES) of tumor and blood normal (BN). **e)** Genomic landscape of the sample cohort showing the top significantly mutated genes. Color scale in heatmap denotes variant allele fraction (VAF) for each gene. All mutations are somatic, unless indicated by a colored circle/triangle/pentagon designating germline variants of different annotated significance. **f)** UMAP plots of all cell types for single-nucleus RNA data colored by cell types. **g)** UMAP plots of all cell types for single-cell RNA data colored by cell types. **h)** UMAP plots of all cell types for single-nucleus ATAC data colored by cell types.

In the normal breast duct, distinct transcriptional programs assign cells to a luminal or basal fate. The regulatory network of *GATA3, FOXA1,* and the estrogen receptor *ESR1* represent a signaling axis that is essential for the maturation of luminal breast cells and the development of luminal BC^20–23^. The ETS-domain transcription factor (TF) ELF5 is a key determinant of luminal cell fate and the secretory sub-lineage of luminal cells^24,25^. Basal breast cells, on the other hand, maintain a more mesenchymal state, with p63 and SOX family TFs playing key roles in the maintenance of basal cell fate^26,27^. Transcriptional programs and chromatin accessibility patterns are useful as markers of cell lineage and cell of origin^28–31^. The structure and chromatin accessibility of mouse breast tissue has established distinct features of cell states and underscored the utility of chromatin accessibility as a marker of breast cell lineage ^32,33^. Chromatin accessibility has also identified master regulators such as SOX10 which regulate the transition from benign breast duct to cancer ^33^. Single-cell chromatin analysis of mammary glands in developing mice have further refined the chromatin signatures associated with normal breast cellular lineages ^34^. Beyond understanding of chromatin accessibility in animal models^34–36^ and within immune cells^16^, there has been no investigation of the epigenetic state of human breast cancers across subtypes and their progenitors at single cell resolution.

Genome-wide expression profiling identified five biological BC subtypes^37^, namely luminal A and B, HER2-enriched, basal-like, and normal breast-like, which differ by hormonal receptor status, proliferation, genomic instability, mutational signatures, treatment response, and prognosis^38–42^. It has long been hypothesized that the high degree of heterogeneity in BC is due to different cells of origin within the breast duct. Evidence has mounted that the similarity between basal-like BC and basal/myoepithelial breast cells is superficial, while the cell of lineage for basal-like BC belongs to the luminal lineage. This paradigm was initially supported by early work on BRCA1-deficient mammary cells, where tumors arise from mammary basal cells^43^. Molyneux and colleagues analyzed a conditional mouse model of *BRCA1* deficiency that developed tumors resembling human basal-like BC, and showed that these arose from a luminal ER-negative (ER-) progenitor population^44^. This is in line with evidence in humans, where *BRCA1* mutation carriers have been shown to harbor an expanded population of luminal progenitor cells with an aberrant phenotype, including expression of some basal epithelial cell markers^45,46^. Gene expression profiling from *BRCA1* heterozygous breast tissue showed similarities between luminal progenitor cells and basal-like breast tumors, and between luminal mature cells and luminal A/B tumors^45^. To address cell of origin, Keller and colleagues isolated luminal (EPCAM^+^CD10^-^) and basal (CD10^+^) cells from *BRCA1* wild-type breast reduction specimens and upon implantation into immunodeficient mice, luminal cells gave rise to tumors resembling both luminal and basal-like subtypes, whereas basal cells gave rise to tumors not closely resembling either basal-like or luminal tumors^47^.

More recently, scRNA-seq gene expression profiling has been used to establish links between breast tumor subtypes and benign duct cell types. Hu and colleagues performed scRNA-seq on breast tumors from *BRCA1* mutation carriers and noncarriers and found similarities between basal-like tumors and the expanded, abnormal luminal progenitor population seen in *BRCA1* carriers, and between ER-positive (ER+) breast tumors and luminal mature cells^9^. Additionally, scRNA-seq from FACS-sorted luminal epithelial cells from reduction mammoplasties showed gene expression similarity between ductal KRT15^+^ luminal progenitors and published signatures of basal-like BC^48^. These studies have largely focused on protein markers and gene expression patterns. In this study, we apply sn/scRNA-seq and snATAC-seq in tandem to clarify not only the gene expression similarities between BC subtypes and their proposed cells of origin, but also the transcriptional networks responsible for transformation and cell lineage identity.

This study aims to understand tumor heterogeneity and its relation to BC lineage at single-cell resolution. Chromatin accessibility, TF motif enrichment, and their impact on the transcriptome reveal the structure of BC heterogeneity through integration of bulk RNA/DNA sequencing, single-cell and single-nucleus RNA sequencing, and single-nucleus ATAC sequencing technologies. Specifically, in this work we explore the transcriptional programs and chromatin accessibility patterns that link breast cancer subtypes to distinct cell types in the benign breast duct. As part of the Washington University Human Tumor Atlas Network (WU-HTAN) program, we generated multi-omic data for 70 samples from 38 ER+PR-HER2-, ER+PR+HER2-, HER2+, and triple-negative breast cancer (TNBC) tumors,4 normal adjacent tissues (NATs) and 1 metastatic liver sample. Of these patients, 27 were treatment-naïve and 12 had undergone previous therapy. We observed subtype-specific chromatin accessibility features associated with driver gene expression signatures. We identified gene expression and chromatin accessibility networks shared between BC subtypes and benign breast duct populations at the single-cell level, which are mapped to specific structures by co-detection by indexing (CODEX) multiplex imaging. These findings may guide our understanding of the early pathogenesis of breast cancer.

## RESULTS

### Clinical features and genomic characterization

We conducted sc and/or sn RNA-seq for 57 tissue samples across 37 resected breast tumors (i.e., “cases”) (**Supplementary Table 1**). Of these, 6, 16, 4, and 11 tumor(s) were clinically annotated as ER+PR-HER2-, ER+PR+HER2-, HER2+, and TNBC, respectively. For a subset of tumors (n = 14), we collected up to three spatially distinct samples from the same tumor using our grid processing method for sample collection (**Fig. 1b-c**). In addition, each sample also underwent extensive imaging characterization and bulk omics. The data generated include scRNA-seq, snRNA-seq, snATAC-seq, spatial transcriptomics (ST), bulk whole exome sequencing (WES), and bulk RNA sequencing (RNA-seq). We generated scRNA data for 31 samples (from 14 cases), snRNA data for 30 samples (from 27 cases), and snATAC data for 26 samples (from 24 cases), of which 4 samples had both scRNA and snRNA for comparison. Additional validation was provided from spatial transcriptomics (ST) data comprising 33 slides from 13 breast cancer cases and CODEX multiplex imaging on 47 slides from 13 cases. 54 and 52 paired samples underwent bulk WES and RNA-seq, respectively.

Of 37 patients with resected breast tumors, samples were obtained from 26 patients pretreatment, and 11 patients following therapy (**Supplementary Table 1**). Systemic treatment regimens for previously treated patients included carboplatin and paclitaxel; doxorubicin and cyclophosphamide followed by paclitaxel; paclitaxel, trastuzumab and pertuzumab; single agent paclitaxel; doxorubicin, cyclophosphamide and pembrolizumab; and aromatase inhibitors. Two patients who had not yet received treatment for the breast tumor included in this study had previously received treatment for prior unrelated breast cancer. The median age of patients was 61. Three patients under the age of 40 were included in this cohort (ages 30, 31, and 38) (**Fig. 1d** **and Supplementary Table 1**). Of these, the patient aged 30 (HT163B1), had a family history including two other family members with breast cancer diagnosed in their thirties. In all, 21 of the 37 patients had a known first-degree family member with a cancer diagnosis, though only 7 of these were known to be breast cancers. The majority of tumors (30 of 37) were histologically identified as invasive ductal carcinoma (IDC) and the other 7 of 37 were invasive lobular carcinoma (ILC) (**Supplementary Table 1**).

We determined somatic and germline variants in the cohort (**Fig. 1e** **and Supplementary Table 2 and Supplementary Table 3**) using WES. Consistent with previous studies, we detected several cases with somatic mutations in *TP53* and *PIK3CA* (**Fig. 1e**). For germline variants, we identified 2 potential pathogenic germline variants in *BRCA2* (p.A938fs in HT243B1, and p.K2013* in HT271B1) and one in *BRIP1* (p.K703fs in HT077B1) using the CharGer pipeline^49^. Notably, these predicted pathogenic germline variants seem to be present at a much higher variant allele fraction (VAF) in the tumor samples compared to paired normals for the affected cases, showing significant loss of heterozygosity (one-sided Fisher exact test adjusted p-values (FDR): 4.60010^-5^ and 4.18010^-5^ for the *BRCA2* variants and 1.98010^-14^ for the *BRIP1* variant). One 30 year old patient with a family history of breast cancer (HT163B1) has a germline frameshift variant of unknown significance (p.Y1672fs) in *BRCA2* that has significant LOH in the tumor (FDR=0.0001). Across spatially separate samples from the same case, we generally detected the same somatic mutations across samples, with a few exceptions likely due to tumor purity. The two cases with somatic mutations in *CDH1* were of lobular histology.

After filtering and quality control (QC), we obtained a total of 138,835 cells and 205,120 nuclei, which we clustered and classified into cell types based on marker gene expression (Methods). For cases with paired WES, we identified copy number alterations that overlap InferCNV calls derived from the single-cell data to confidently identify tumor subpopulations relative to normal cells (**Extended Data Fig. 1a and Supplementary Table 4**). In addition to tumor cells, we identified stromal cells of the breast, including endothelial cells, cancer-associated fibroblasts of the vascular (vCAF), matrix (mCAF), developmental (dCAF) and cycling (cCAF) subsets, and adipocytes. Within the benign breast compartment, we captured benign duct cells including luminal mature cells, luminal progenitor cells, and basal/myoepithelial cells. Lymphocyte subsets include B cells, plasma cells, and CD4+ or CD8+ T cells, with T cells being further subdivided into including regulatory (Treg), cytotoxic, pre-exhausted, exhausted, activated, and proliferating subsets. Other immune components including monocytes, macrophages, dendritic cells including classical (cDC1, cDC2) and plasmacytoid (pDC), NK cells, NKT cells, and mast cells were also identified (**Fig. 1f-h**). We calculated the fraction of tumor cells for samples with adequate coverage ranging from 1.6% to 82% in scRNA and 1.5% to 99% in snRNA. Related to other work, this study provides high quality single-cell data (snRNA: mean 2187 genes/cell; scRNA: mean 2448 genes/cell) relative to other large cohorts for breast cancer ^11^ (**Extended Data Fig. 1b**). As previously reported, while snRNA-seq and scRNA-seq both capture similar cell type composition in each assay, the proportions can vary dramatically with frozen tissue nuclei isolation techniques (snRNA) capturing a higher tumor fraction and fresh tissue whole cell (scRNA) dissociation capturing more immune cells^50,51^. To take advantage of these differences, we explored tumor heterogeneity using snRNA-seq/snATAC-seq and the tumor microenvironment using scRNA-seq. Further, for some cases with paired scRNA-seq and snRNA-seq data (from different regions of the same tumor), we validated findings using the orthogonal method. In summary, we have generated a large compendium of single-cell data encompassing both RNA (snRNA-seq and scRNA-seq) and ATAC data spanning three subtypes of breast cancer and normal adjacent tissues to study tumor heterogeneity and normal to tumor transition states.

### Tumor subtype intrinsic and extrinsic characterization

Historically, breast tumor subtype assignments are calculated from bulk RNA-seq data using published methods for the PAM50 assay^52^. However, assigning tumor subtype from gene expression is confounded by the composition of tumor and non-tumor cells in a sample. To disentangle subtype assignment from stromal contribution, the PAM50 algorithm was applied separately to bulk RNA-seq and snRNA-seq data (Methods). Subtype assignments from bulk RNA-seq and snRNA-seq demonstrate good concordance: 12 of 14 samples (85%) with both bulk RNA-seq and snRNA-seq had identical PAM50 calls from both modalities. Of the discrepant cases, the bulk RNA-seq based assignments were normal-like and luminal A, and both cases were called luminal B from snRNA-seq. PAM50 subtype assignments from our cohort (**Supplementary Table 1**) also closely mirrored clinical biomarker profiles. Thirteen of 15 TNBC samples with bulk RNA-seq or snRNA-seq data (87%) were assigned to the basal-like subtype, with two assigned as HER2-enriched (**Fig. 1d** **and Supplementary Table 1**). Sixteen of 38 (42%) clinically defined ER-positive, HER2-negative samples (with or without PR positivity) with bulk RNA-seq or snRNA-seq data were assigned to the luminal A subtype, with another 19 (50%) assigned to the luminal B subtype. The remaining four (8%) ER-positive, HER2-negative samples were assigned to the basal-like subtype. Three clinically HER2-positive samples were included in this data set, and all were classified as HER2-enriched by PAM50.

Just as average expression of PAM50 genes in tumor cells from snRNA-seq could discriminate tumor subtype, chromatin accessibility in the promoters of PAM50 genes from snATAC-seq showed good segregation of tumors by subtype in the 21 samples (from 21 cases) with both data types available (**Fig. 2a-c**). In addition to BC cells, benign epithelial ductal cells were identified and stratified into three benign cell types, using both published expression markers and co-clustering status across samples ^6,7,9^. Each benign cell type harbored unique markers, namely *KIT*, *KRT15*, and *PTPN* for luminal progenitor cells (LP)*, ANKRD30A, ERBB4*, and *AFF3* in luminal mature (LM), and *ACTA2*, *RBMS3,* and *DST* in basal/myoepithelial progenitor (BP) cells. We did not identify a robust population of mammary stem cells, consistent with their low abundance in adults^4^. Benign ductal cells were detected in all clinical subtypes (ER+, ER+/PR+, HER2+, and TNBC) and PAM50 subtypes (luminal A, luminal B, HER2-enriched, and basal-like) (**Extended Data Fig. 1c)**. Across all samples, we identified all three progenitor cell types in 46% (n = 24) and 36% (n = 14) of samples in the scRNA and snRNA cohorts, respectively (**Extended Data Fig. 1d**). Compared to other benign ductal cells, luminal mature cells expressed high levels of *ERBB4, DACH1, and ESR1*, with hormone-response pathways enriched among differentially expressed genes (**Fig. 2b** **and Supplementary Table 5**). In contrast, luminal progenitor cells were characterized by high *KIT* expression as well as expression of other progenitor markers (*ALDH1A3*). Finally, basal/myoepithelial cells showed high expression of genes involved in cytoskeleton and myoepithelial contraction, including *ACTA2* and *DST*, as well as *TP63*. (**Fig. 2b**). Genes included in the PAM50 subtyping assay show dramatic differences by subtype even by snRNA measurements and are further confirmed in the snATAC data (**Fig. 2c**). Differentially accessible promoters by subtype highlighted key subtype-associated genes including *VIM* and *SOX4* in basal-like tumors, *FOXA1* and *GATA3* in luminal tumors, and *ERBB2* and *GRB7* in HER2-enriched tumors (**Fig. 2d**). Promoter accessibility of PAM50 genes showed stark subtype differences and highlighted similarities to benign duct populations (**Fig. 2e**). Key basal-like genes *SFRP1* and *KRT17* showed high promoter accessibility in basal-like tumors and luminal progenitor cells, while key luminal genes *ESR1* showed promoter accessibility in luminal A/B tumors and luminal mature cells. By analyzing over twenty samples from various patients, we have built a large resource of both breast cancer cells and benign duct populations, enabling us to evaluate the transcriptional programming responsible for the normal to tumor cell transition across multiple subtypes of breast cancer.

**Figure 2.**
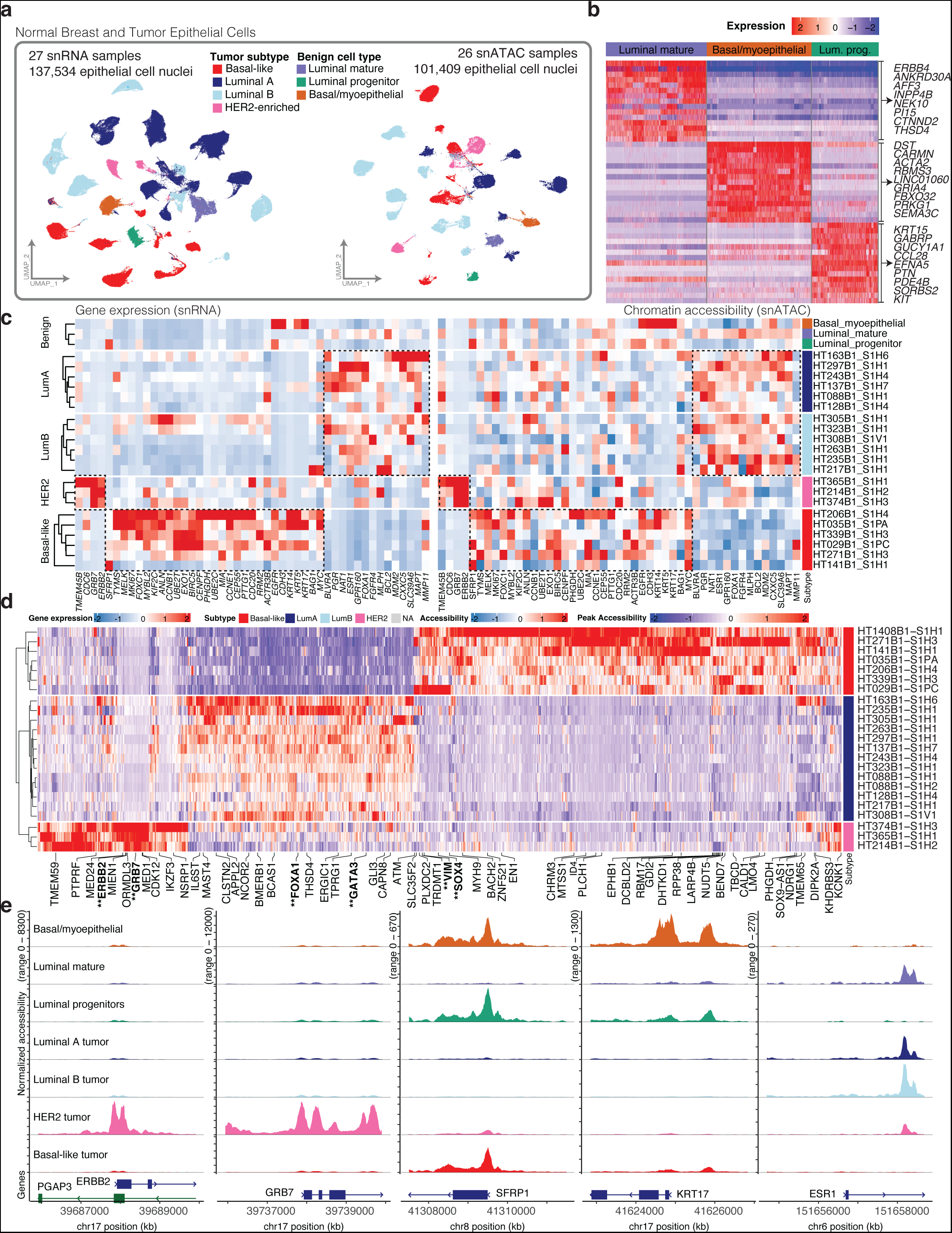
Tumor Subtype and Benign Duct Cell Types. **a)** UMAP plots of benign breast epithelial cells and breast cancer cells for all snRNA (left) and snATAC (right) samples. Tumor cells colored by PAM50 subtype. **b)** Heatmap of top 15 differentially-expressed genes in snRNA-seq data from benign breast duct cells. A subset of genes from each benign cell type is highlighted in the figure. **c)** Heatmaps of snRNA gene expression (left) and snATAC chromatin accessibility (right) for genes in the PAM50 subtyping assay. Average values are shown for all tumor cells per sample, as well as each benign breast duct cell type pooled across samples (top). Characteristic genes identifying luminal A/B, HER2-enriched, and basal-like subtypes are shown in boxes. **d)** Peak accessibility for differentially accessible promoters by breast cancer subtype in snATAC-seq data. Key subtype-associated genes are highlighted in bold and with two asterisks below. **e)** Coverage plots showing normalized chromatin accessibility across promoter regions of key subtype-associated genes in snATAC-seq data from tumor nuclei grouped by subtype, and benign epithelial cell types.

In addition to profiling tumor and benign ductal cells, we also examined subtype differences in the immune compartment. Lymphoid and myeloid cells were profiled in 31 scRNA-seq samples comprising 29 tumor samples and two NAT samples (**Fig. 3a**). Exhausted CD8+ T cells were significantly more prevalent in basal-like tumors compared to luminal A or B tumors (**Fig. 3a-b**). This finding was consistent in snRNA-seq data and independent of treatment (**Fig. 3b** **and Extended Data Fig. 2a-b**). We performed cell-cell interaction analysis using CellPhoneDB and observed significant predicted interactions between *CTLA4* expressed by CD8+ T cells and CD86 expressed by multiple myeloid cell types (macrophages, monocytes, cDC1, and cDC2) in basal-like samples relative to luminal samples (**Fig. 3c**) (Methods)^53^. *CTLA4* on CD8+ T cells was also predicted to interact with *CD80* on various myeloid cell types in basal-like tumors, though this did not reach statistical significance. Compared to luminal A or B tumors, exhausted CD8+ T cells in basal-like tumors expressed higher levels of *CTLA4*, *CXCL13*, and *CCL3* (**Extended Data Fig. 2c)**. To validate this finding, we utilized spatial transcriptomics data from the 10x visium platform. For two basal (HT206B1, HT271B1) and two luminal samples (HT323B1, HT262B1), the ST spots overlapping lymphocyte dense regions were extracted (**Fig. 3d** **and Extended Data Fig. 3**). We hypothesized that if cell-cell interactions between myeloid and T-cell populations maintained differential interactions between subtypes as shown in the single-nuclei data, then this would hold true for each subtype in lymphocyte dense regions derived from ST data. The ST data confirmed that *CTLA4*, *CD80*, *CD86* and *CD1C* had an overall higher expression across 2 basal samples relative to the 2 luminal samples (**Fig. 3d**). Finally we performed cell type deconvolution using CytoSPACE on 33 slides from 4 basal-like, 8 luminal and 1 HER2 breast cancers and again observed increased abundance of exhausted CD8+ T cells in basal-like cancers (**Fig. 3b** **and Extended Data Fig. 4**)^54^.Taken together, this provides evidence of increased immunosuppression and exhaustion in T cells in basal-like breast tumors.

**Figure 3.**
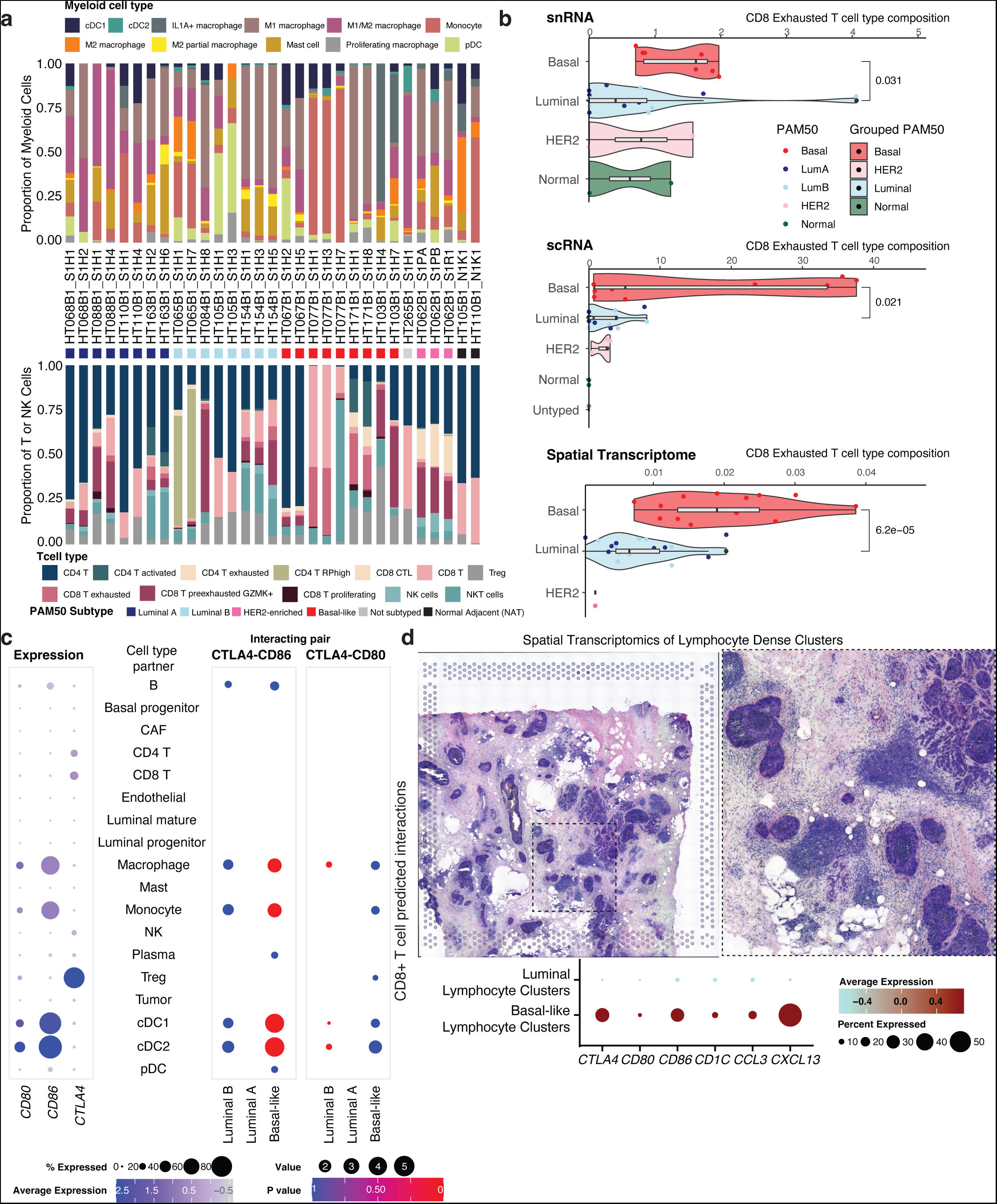
Subtype-enriched Elements of the Tumor Microenvironment. **a)** Composition of myeloid immune subsets (top) and T/NK subsets (bottom) for each sample with single-cell RNA-sequencing data. **b)** Proportion of Exhausted CD8 T cells by subtype identified by snRNA seq, scRNA seq and Spatial Transcriptomics (ST). Each dot is the proportion of exhausted CD8 T cells relative to other T cells for an individual piece for the snRNA and scRNA while for the ST it is based on the proportion of total spots. **c)** Left plot indicates the expression of three markers (*CD80*, *CD86*, and *CTLA4*) in the RNA. Size of dot indicates % of genes expressing the gene and color indicates average expression. Right plot summarizes CellPhoneDB results indicating interacting gene partners in the single-cell RNA-seq data. Size of dot indicates mean expression of interacting gene partners in their respective cell types and color indicates p-value. **d)** Top images show an example of a lymphocyte dense region in one sample of interest. Right image is a zoomed-in region of the left image which we use to quantify the expression of various markers in the lower panel. Expression of a subset of genes in lymphocyte dense clusters isolated from spatial transcriptomics data from luminal and basal cancers. Size of dot indicates percent of the spots included in the analysis that express the gene of interest while color indicates average expression.

### Cell of origin and transcriptional regulons of putative tumor lineages

Identifying and understanding the cells that give rise to BC is critical for comparing tumor and normal cells and ultimately for understanding tumor progression and evolution. While there is no consensus in the field of which precise cell types give rise to tumor cells in breast cancer, the prevailing model is that luminal progenitor cells tend to give rise to TNBC cancers, while luminal mature cells develop into ER+ or ER+PR+ tumors^4,47^. The chromatin landscape at single-cell resolution is uniquely suited to reconstruct the lineage between progenitor populations and malignant cells in a tumor. To determine whether tumor subtypes were associated with distinct cells of origin, we performed Monocle trajectory analysis on snATAC-seq data from basal-like and luminal A/B tumor cells and benign luminal mature, luminal progenitor, and basal/myoepithelial cell populations (Methods). We observed for the majority of basal-like cases, that tumor cells were closely associated with luminal progenitor cell populations, while for the majority of the luminal cases, we observed tumor cells to be closer to luminal mature cells (**Fig. 4a-b**). Correlation of motif scores across epithelial cell types in individual cases also highlighted greater similarity between basal-like BC and luminal progenitor cells and between luminal BC and luminal mature cells (**Fig. 4c-d**). Finally, motifs showing high chromatin accessibility in luminal progenitor cells were also highly represented in open chromatin in basal-like breast tumors (**Fig. 4e**), while motifs found in luminal mature cells were also highly represented in open chromatin in luminal breast tumors (**Fig. 4f**). Motifs that exhibited differential accessibility in luminal mature cells and were also enriched in open chromatin in luminal tumors included forkhead family proteins, GATA3, ESR1, and HNF1A. Differentially accessible motifs for luminal progenitor cells also enriched in open chromatin in basal-like tumors included GRHL1 and TFCP2. The TFs for which accessibility was correlated with pseudotime between luminal tumors and luminal mature cells, and between basal-like tumors and luminal progenitor cells, are shown in Extended Data Fig. 5.

**Figure 4.**
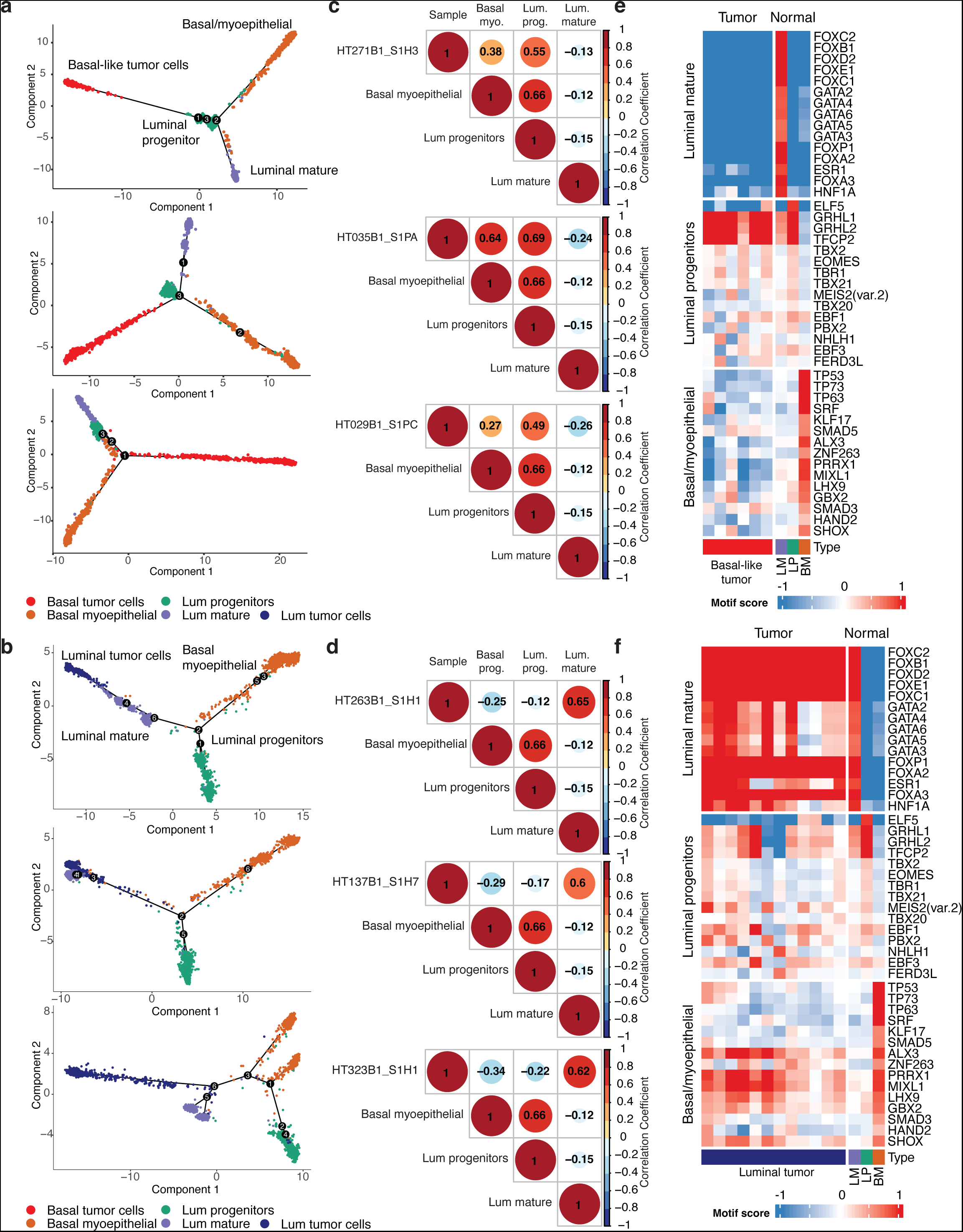
Chromatin Accessibility Evidence for Subtype-specific Cell of Origin. **a)** Monocle pseudotime plots of tumor and benign breast duct cells from three representative basal-like breast cancer samples. **b)** Monocle pseudotime plots of tumor and benign breast duct cells from three representative luminal breast cancer samples. **c)** Correlation matrices for transcription factor (TF) motif scores from tumor cells and benign duct cells for the breast cancer samples in (A). **d)** Correlation matrices for TF motif scores from tumor cells and benign duct cells for the breast cancer samples in (B). **e)** Heatmap of motif scores for the top 15 differentially accessible motifs identified in luminal mature (LM), luminal progenitor (LP), and basal/myoepithelial (BP) cells. Scores are shown for tumor cells from each basal-like snATAC-seq sample and for benign breast duct cells. **f)** Heatmap of motif scores for the top 15 differentially accessible motifs identified in luminal mature, luminal progenitor, and basal/myoepithelial cells. Scores are shown for tumor cells from each luminal snATAC-seq sample and for benign breast duct cells.

To explore pre-cancer states during early malignancy, we evaluated tumor and normal cells in the MMTV-PyVT mouse model of luminal breast cancer^55^. Mouse mammary glands were collected at 12 weeks old to capture the transition of normal ducts to cancer cells. Both normal ducts and cancer cells were recognized in the H&E staining of embedded samples (**Extended Data Fig. 6a-b**). This is concordant with snATAC data derived at the same time point showing that both early stages of cancer cells and normal ducts are present in the mouse model. Trajectory analysis using monocle on the ATAC data again confirmed the transition of proposed cancer cells (Lum_0, Lum_2, Lum_4, Lum_6) from luminal mature cells rather than progenitors or basal/myoepithelial cells (**Extended Data Fig. 6c-d**).

To confirm markers for single-cell/single-nucleus populations and provide support for benign cell types and their connection to putative cell of origin, we performed CODEX multiplex imaging on representative basal-like and luminal BC sections. SMA, podoplanin, and vimentin were used for staining basal myoepithelial cells, while c-Kit and GATA3 were used for luminal progenitor cells and luminal mature cells, respectively (Methods). As a result, tumor cells from the luminal B tumor HT323B1 exhibit tumor cells positive for ER and PR, with lower proliferative signature by Ki67 staining and a lack of c-Kit staining (**Fig. 5a**). Tumor cells from basal-like tumors had a strong proliferative Ki67 staining, and high c-Kit positivity (**Fig. 5b**). From the benign structures, basal myoepithelial cells are positive for SMA, podoplanin, and vimentin, luminal mature cells are positive for GATA3, and luminal progenitor cells are positive for c-Kit (**Fig. 5c**). Notably, normal duct cells coexpressing c-Kit and GATA3 were rare and may have been due to cell segmentation errors. Quantification of immunofluorescence signal in areas of tumor and normal duct showed higher c-KIT positivity in normal duct compared to luminal tumor, but higher c-KIT positivity in basal tumor compared to normal duct (**Fig. 5e**). The increased c-KIT positivity in both basal tumor regions and normal luminal progenitors further emphasizes the connection between these two cells from a proteomic view. Epithelial cell markers in normal duct and tumor populations in basal-like and luminal tumors are shown in Fig. 5e. Similarly, GATA3 showed increased positivity in normal duct and luminal tumor and decreased positivity in basal tumors (**Fig. 5e**). Additional epithelial markers were quantitated in tumor and normal duct regions with expected results, including increased CK14 positivity in basal tumors and increased ER and PR positivity in luminal tumors (**Fig. 5e**). Gene expression and chromatin accessibility for CODEX marker genes in snRNA-seq and snATAC-seq are consistent with these results (**Fig. 5 f-g**).

**Figure 5.**
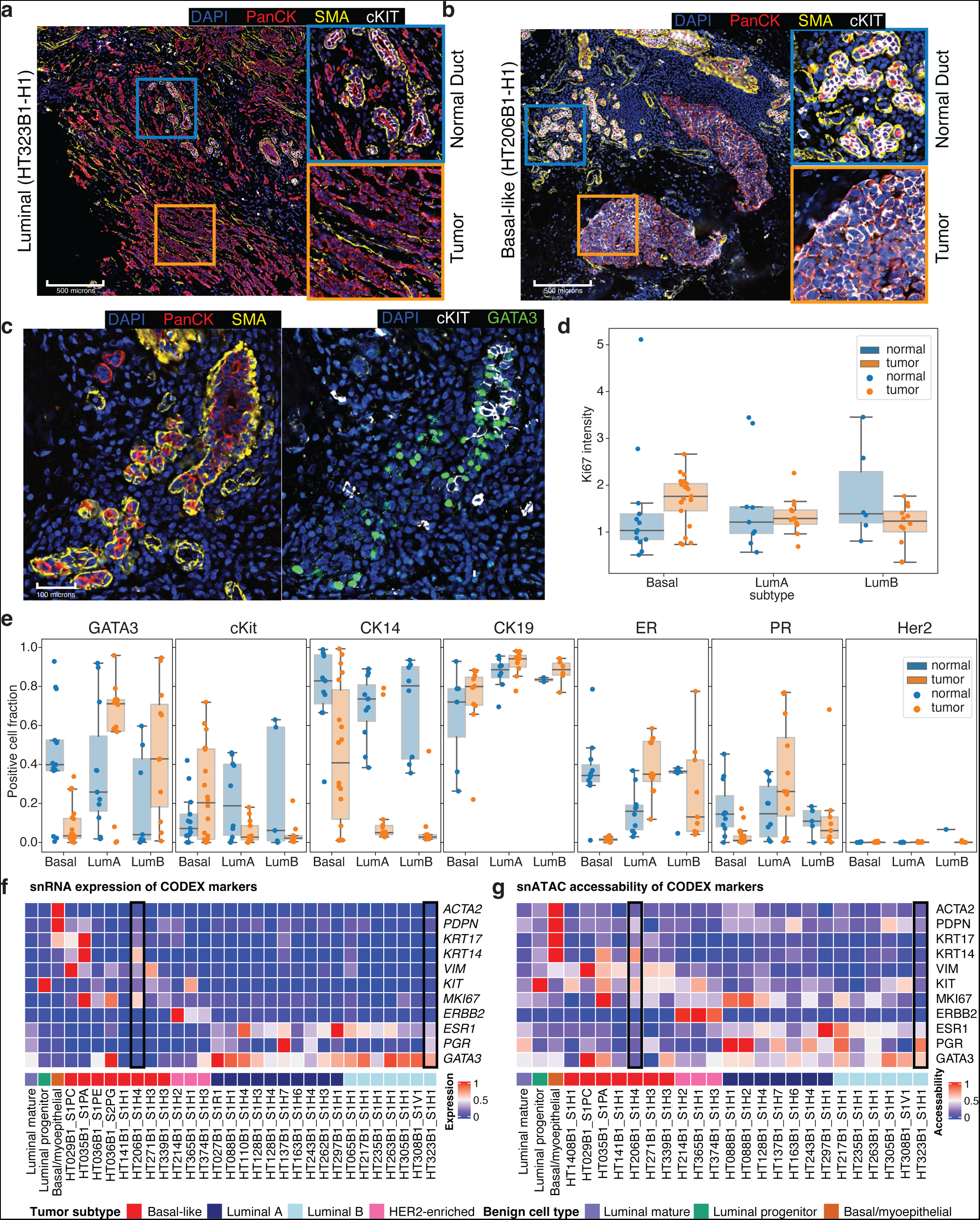
Spatial Characterization of Tumor Subtype and Normal Ducts. **a)** CODEX multiplex immunofluorescence on luminal sample HT323B1. Inset regions (square) are expanded to the right and colored by related inset. DAPI is stained in blue, PanCK in red, SMA in yellow and cKIT in white. **b)** CODEX multiplex immunofluorescence on basal sample HT206B1. Inset regions (squares) are expanded to the right and colored by related inset. DAPI is stained in blue, PanCK in red, SMA in yellow and cKIT in white. **c)** Section of CODEX immunofluorescence image from HT206B1 centered on a benign ductal region. Section on the left is stained with DAPI in blue, PanCK in red, and SMA in yellow. The section on the right is stained with DAPI in blue, cKIT in white, and GATA3 in green. **d)** Boxplot summarizing overall Ki67 intensity across all samples in normal duct and tumor regions separated by subtype. **e)** Positive cell fraction of GATA3, cKit, CD14, CK19, ER, PR and Her2 across all samples in normal duct and tumor regions separated by subtype. **f)** Average expression scores of CODEX marker genes in the single-nuclei RNA sequencing data. Gene expression for samples HT206B1_S1H1 and HT323B1_S1H1 used for CODEX imaging are outlined. **g)** Average chromatin accessibility scores of CODEX marker genes in the single-nuclei ATAC sequencing data. Chromatin accessibility for samples HT206B1_S1H1 and HT323B1_S1H1 used for CODEX imaging are outlined.

To explore tumor cell of origin more deeply, we sought to reconstruct transcriptional networks specific to these distinct lineages. Grouping differentially accessible motifs of tumor cells from snATAC-seq, a high degree of similarity between luminal progenitor cells and basal-like tumor cells and between luminal mature cells and luminal A/B tumor cells was observed, while basal myoepithelial cells were distinct from all tumors (**Fig. 6a**). Key TF motifs enriched in the open chromatin of basal-like tumors and luminal progenitor cells include NFIB, TEAD family TFs, SOX family TFs, and CEBPB. In contrast, luminal A/B tumor cells and luminal mature cells showed high accessibility for the estrogen receptor ESR1, as well as forkhead proteins including FOXA2 and FOXP1, GATA3 and other GATA-box transcription factors, and HNF1A. Two HER2-enriched samples had snATAC-seq data and showed enrichment for RARA, NR6A1, and ESRRA motifs. Basal myoepithelial cells showed high motif accessibility for TFs including TP63, ZBTB18, and SRF.

**Figure 6.**
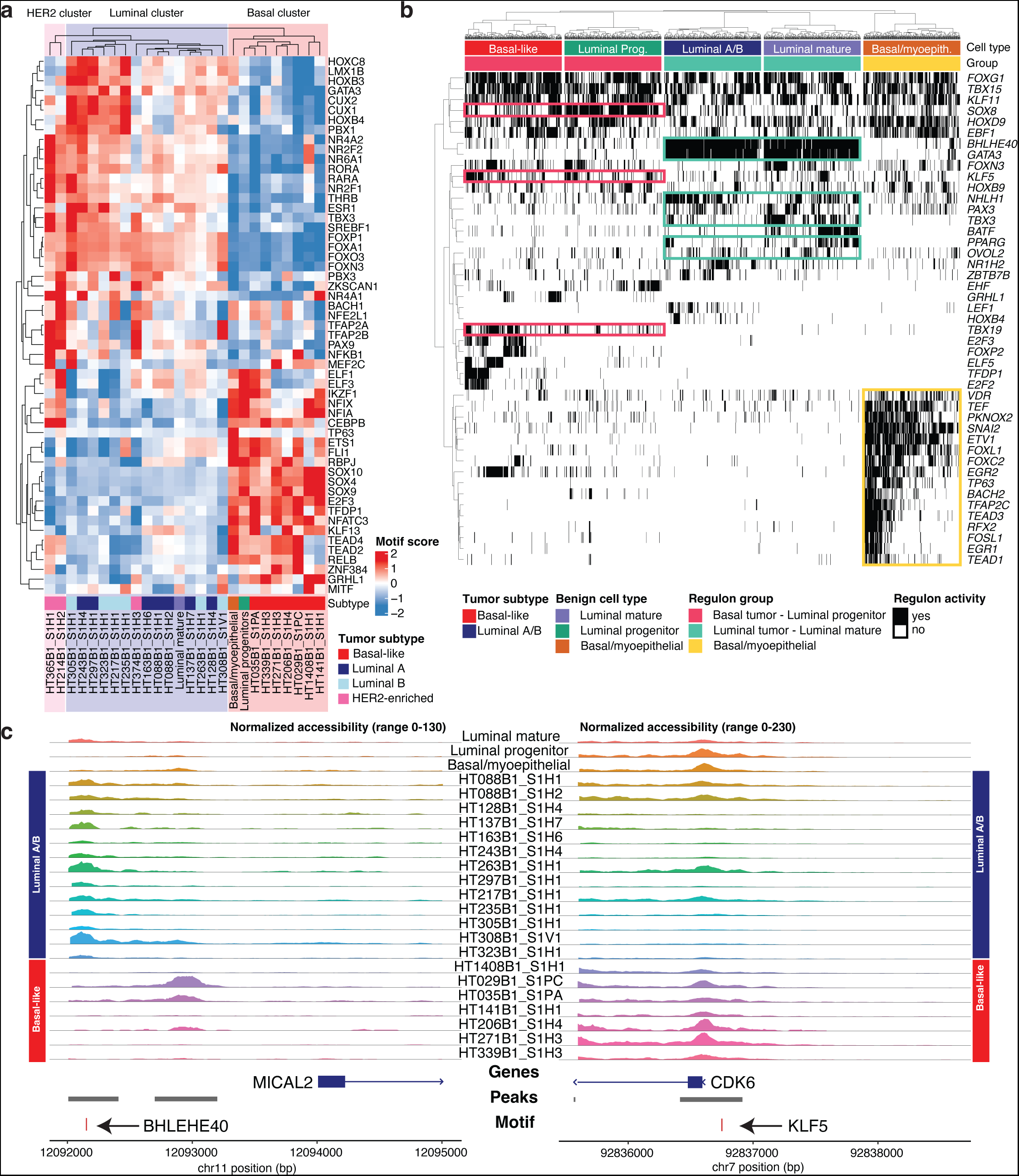
Tumor Lineage-specific Regulators of Gene Expression. **a)** Heatmap of differentially accessible motifs identified in tumor cell snATAC-seq data. Motif scores are shown for average value across tumor cells in each sample and for luminal progenitor, luminal mature, and basal/myoepithelial cells pooled across all samples. **b)** Binarized heatmap of regulon activity in tumor-normal lineage groups. Color bars above show tumor/benign cell type and regulon group (basal-like BC and luminal progenitor, luminal A/B BC and luminal mature, and basal myoepithelial). **c)** Coverage plots of normalized snATAC-seq accessibility across promoter regions for *MICAL2* (left) and *CDK6* (right). Regulon transcription factor motifs and ATAC peak regions shown below.

To interrogate the genes linking tumor cells and their proposed cell of origin, we next used SCENIC to identify regulons of co-regulated genes in snRNA-seq data (Methods)^56^. Based on the evidence from motif accessibility and pseudotemporal association (**Fig. 4**), regulons were identified for three related lineage groups: (1) basal-like tumor and luminal progenitor cells, (2) luminal tumors and luminal mature cells, and (3) basal myoepithelial cells. Lineage-specific regulons for lineage group 1 include SOX8, KLF5, and TBX19 (**Fig. 6b**). In contrast, lineage-specific regulons identified for group 2 include BHLHE40, GATA3, NHLH1, PAX3, TBX3, OVOL2, and PPARG. Finally, regulons specific to basal myoepithelial cells include VDR, SNAI2, ETV1, and TP53. Of note, while *SOX8* expression has been implicated in TNBCs as a regulator of stem-like capabilities in tumor cells, its expression in luminal progenitors has not been described^57^. Within lineage group 2 (luminal tumors and luminal mature cells), the co-regulation of *BHLHE40* and *GATA3* are likely the result of hypomethylation in luminal A tumors^58^, which is also shared by luminal mature cells in our dataset.

To provide orthogonal evidence for the importance of these TFs, we sought genes in these lineage-specific regulons with the regulon TF motif located in a differentially accessible ATAC peak in the gene’s promoter. Examples of this include the BHLHE40 motif in the promoter region of *MICAL2* and the KLF5 motif in the promoter region of *CDK6* (**Fig. 6c**). Within our sample cohort, the BHLHE40 motif upstream of *MICAL2* is accessible in luminal mature and luminal A/B tumor cells and is less accessible in basal-like tumor samples. BHLHE40 is a transcriptional regulator whose overexpression is associated with metastatic potential and malignant proliferation^59^. MICAL2 is involved in modifying the cellular cytoskeleton, and in BC, its overexpression is associated with cell migration via the EGFR signaling pathway^60^. While the expression of *BHLHE40* in luminal tumors has been noted due to hypomethylation^58^ and the expression was reported to increase between normal and invasive tissues^61^, the relationship between BHLHE40 and downstream targets has not been extensively explored between subtypes and within the luminal mature population. More notably the relationship between the transcription factor BHLHE40 and the downstream gene *MICAL2* has not been reported in BC. As noted here, the integration of chromatin accessibility and gene regulation can distinguish the relationships between the progenitor populations and different subtypes of tumor cells and highlights specific transcription factor regulatory networks that define this relationship.

### Lineage specific changes along the spectrum from progenitor to tumor

To explore lineage specific transcriptional changes between the putative cell of origin and tumor subtypes, we evaluated overlapping and unique differential gene expression profiles for each epithelial cell subset. Using a filtering strategy by comparing differentially expressed genes (DEGs) between related lineages (luminal progenitor and basal tumor, luminal mature and luminal tumors) and removing genes specific to uninvolved subsets (ex. basal myoepithelial, HER2 tumor), we identified 44 genes specific to the basal lineage and 54 genes specific to the luminal lineage (Methods) (**Fig. 7a-b** **and Extended Data Fig. 7 a-b**). Expression of *CCL28*, *APP*, *EHF*, *LINC00342*, among others, is increased in the luminal progenitor relative to the basal tumor. Of note, ETS homologous factor (EHF) is reported to be an anti-EMT factor^62^ and its decreased expression observed in tumor cells supports the finding that basal tumors tend to have increased EMT properties relative to luminal tumors. Basal tumors on the other hand have increased expression of *PRKCA*, *SOX6*, *RGS6*, *CARD18* and several long noncoding RNAs, compared to the progenitor. The role of SOX family members, including *SOX6*, is well documented in basal-like breast cancer^27^. The serine-threonine kinase *PRKCA* has been observed to be upregulated in breast cancer, inversely correlated with estrogen receptor expression, and a critical member of signaling networks in cancer stem cells, and thus is being explored as a therapeutic target in TNBC^63^. Finally, several genes share maintained expression between the progenitor and the tumors including *SOX9-AS1*, *GABRP* and *ELF5*. *GABRP* has been observed as an upregulated gene in TNBC and was found to maintain EGFR signaling in BC cell culture and contribute to chemoresistance in BC xenograft models^64^. ELF5 is a transcription factor involved in mammary stem cell fate^65^. Interestingly, for the luminal subtype, we find that many DEGs are shared between the comparisons of luminal A subtype and luminal mature, and between luminal B subtype and luminal mature, but not much overlap between all three groups. This suggests that different mature ductal progenitors may give rise to each luminal subtype. Regardless of luminal subtype (A/B), luminal mature cells had increased expression of *ELOVL5*, *EFHD1*, *NEK10*, *LYPD6* and *NOVA1*, among others, relative to the tumor cells. *ERBB4*, *NOVA1*, and *LINC02306* had relatively stable expression in luminal B and luminal mature cells. Luminal A and luminal B tumors shared increased expression of *FAM155A* and *LRP1B* compared to luminal mature cells, although to a greater extent in luminal B. Within the luminal A subtype, nuclear *LRP1B* was found to correlate with poor prognosis, though the mechanism of its role in carcinogenesis is unclear^66^. Luminal A and luminal mature cells share expression of *PGR*, *THSD4*, *PRLR*, and *ANKRD30A,* which are dramatically decreased in luminal B tumors. For a subset of genes with ATAC gene activity measured by Signac, we were able to validate the increased or decreased accessibility of the luminal and basal lineage genes in the ATAC data (**Extended Data Fig. 7c**).

**Figure 7.**
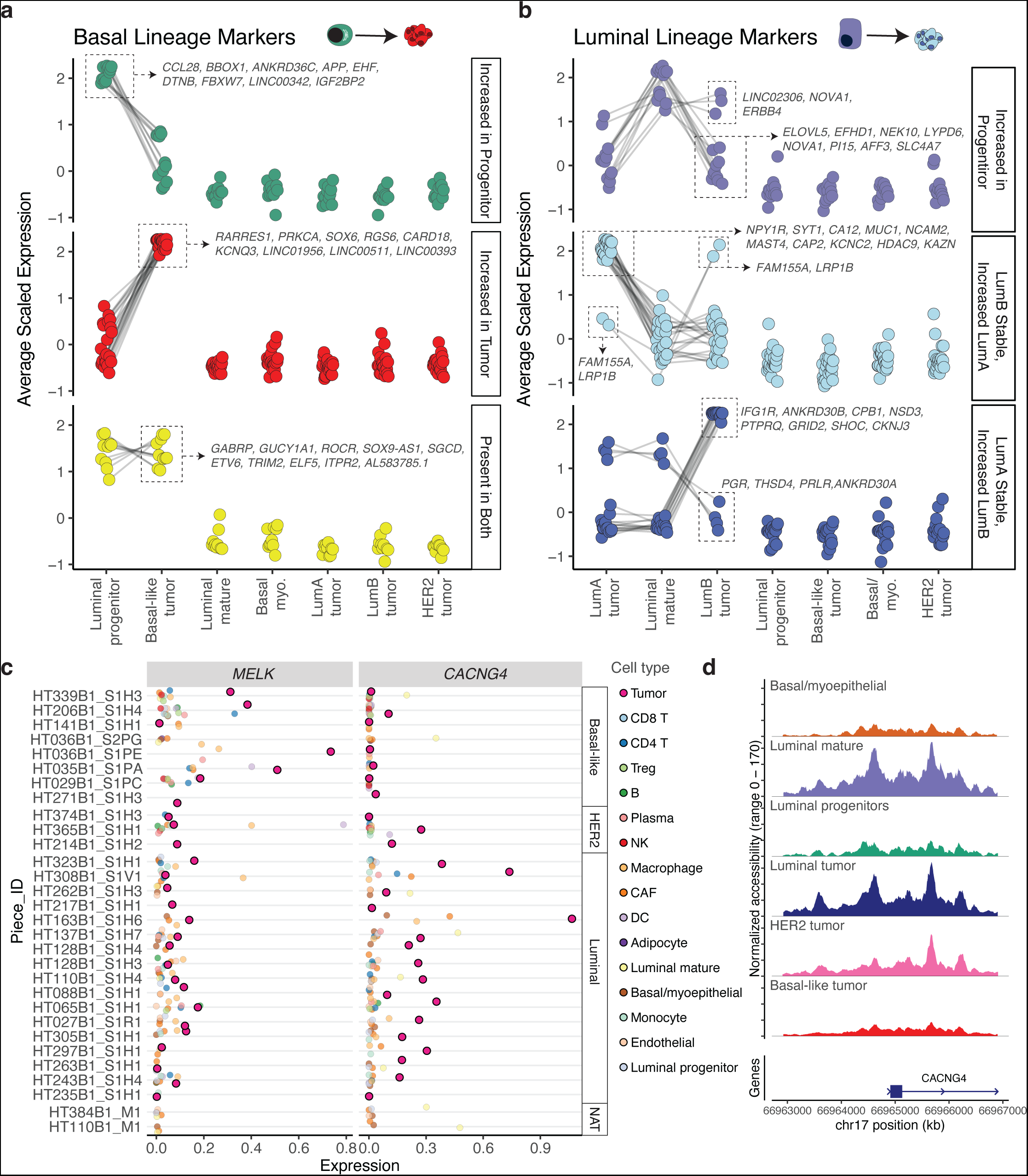
Differential Markers of Basal-like and Luminal Breast Cancer Lineage. **a)** Dot plots showing average scaled expression of basal-like breast cancer lineage markers. Markers are divided into genes expressed highly in luminal progenitor cells but not in basal-like breast cancer (top), genes with increased expression in basal-like breast cancer compared to luminal progenitor cells (middle), and genes high in both groups (bottom). Gene lists are shown for specific groups. **b)** Dotplots showing average scaled expression of luminal A/B breast cancer lineage markers. Markers are divided into genes expressed highly in luminal mature cells but not in luminal BC (top), genes with increased expression in luminal A BC compared to luminal mature cells (middle), and genes with increased expression in luminal B BC compared to luminal mature cells (bottom). Dot size indicates average scaled gene expression. Gene lists are shown for specific groups. **c)** Gene expression across cell types of cell-surface tumor specific markers: *MELK* identified for basal samples, and *CACNG4* identified for luminal samples. **d)** Coverage plot of normalized snATAC-seq chromatin accessibility across the promoter region of *CACNG4* for tumor subtypes and benign breast cell types.

We utilized a similar approach to evaluate differentially accessible motifs (DAMs) to uncover TFs that are important in maintaining lineage identity (*i.e.*, those highly active in related lineages: luminal progenitor and basal tumors, luminal mature and luminal tumors) and TFs that change between benign populations and their related cancer type. We identified 57 TFs enriched in the basal lineage and 47 TFs enriched in the luminal lineage (**Extended Data Fig. 8a-b**). The motifs for TFs GRHL1, GRHL2, TFCP2, and HOXD13 were over-represented in open chromatin regions in both basal BC and LP cells, whereas several TFs including YY1 and YY2, E2F1 and E2F4, SOHLH2, PROX1, OTX1, NFYC, THAP11, ZNF140, and CENPF showed further increase in enrichment in basal BC beyond that seen in luminal progenitor cells. YY1 has been implicated as promoting TNBC via a long noncoding RNA mechanism leading to degradation of PTEN^67^. The role of PROX1 in breast cancer is not well described, but it belongs to a family of genes that drive cell invasion, and PROX1 has been hypothesized to drive invasiveness in colorectal cancer and Kaposi sarcoma^68,69^. Predictably, TFs related to proliferation (including E2F1, E2F4, and CENPF) were increased in basal BC compared to benign luminal progenitor cells. Luminal BC and luminal mature cells showed enrichment of ESR1 and ESR2, GATA family TFs, POU domain TFs, CUX1, CUX2, and PPARG, among others, compared to other benign breast cell types and other BC subtypes. There was overall less divergence between luminal BC and luminal mature cells in terms of TF activity, with nearly all enriched luminal BC motifs also showing enrichment in luminal mature cells.

### Basal-like and luminal tumor cell-surface markers

To identify potential therapeutic targets, we searched for cell-surface tumor specific markers in samples of basal and luminal subtypes (Methods). By this analysis, we initially identified three cell surface genes expressed in basal-like samples, and four in luminal A/B samples. Among those, two cell surface markers were exclusive to either basal or luminal A/B tumors: MELK in basal-like samples, and CACNG4 in luminal A/B samples. MELK is a cell cycle regulator and it is known to be specifically up-regulated in BC samples of basal subtype ^70^ (**Fig. 7c** **and Extended Data Fig. 8c**). CACNG4 is a calcium channel subunit that was previously reported to be associated with metastasis in BC, and it was also found to be highly expressed in ER-positive BC cell lines^71^. Furthermore, we examined promoter accessibility of those markers, and we observed that the promoter region of CACNG4 is more open in luminal tumor and luminal mature cells (**Fig. 7d**). The increased expression of MELK in basal samples was validated with immunofluorescence staining across 4 samples (two basal and two luminal) confirming the increased expression in regions of basal samples relative to luminal (**Extended Data Fig. 9-10**). VTCN1 was also identified as a tumor-enriched cell surface marker, and is clinically relevant as antibody-drug conjugates targeting VTCN1 are currently being evaluated in clinical trials. However, this marker was not subtype-specific. Taken together, we have started to clarify the gene regulatory networks driving the progression of a progenitor cell to luminal and basal-like BC tumor subtypes over the course of tumorigenesis with a focus on expression alterations, motif enrichment, and chromatin accessibility (**Fig. 8a**).

**Figure 8.**
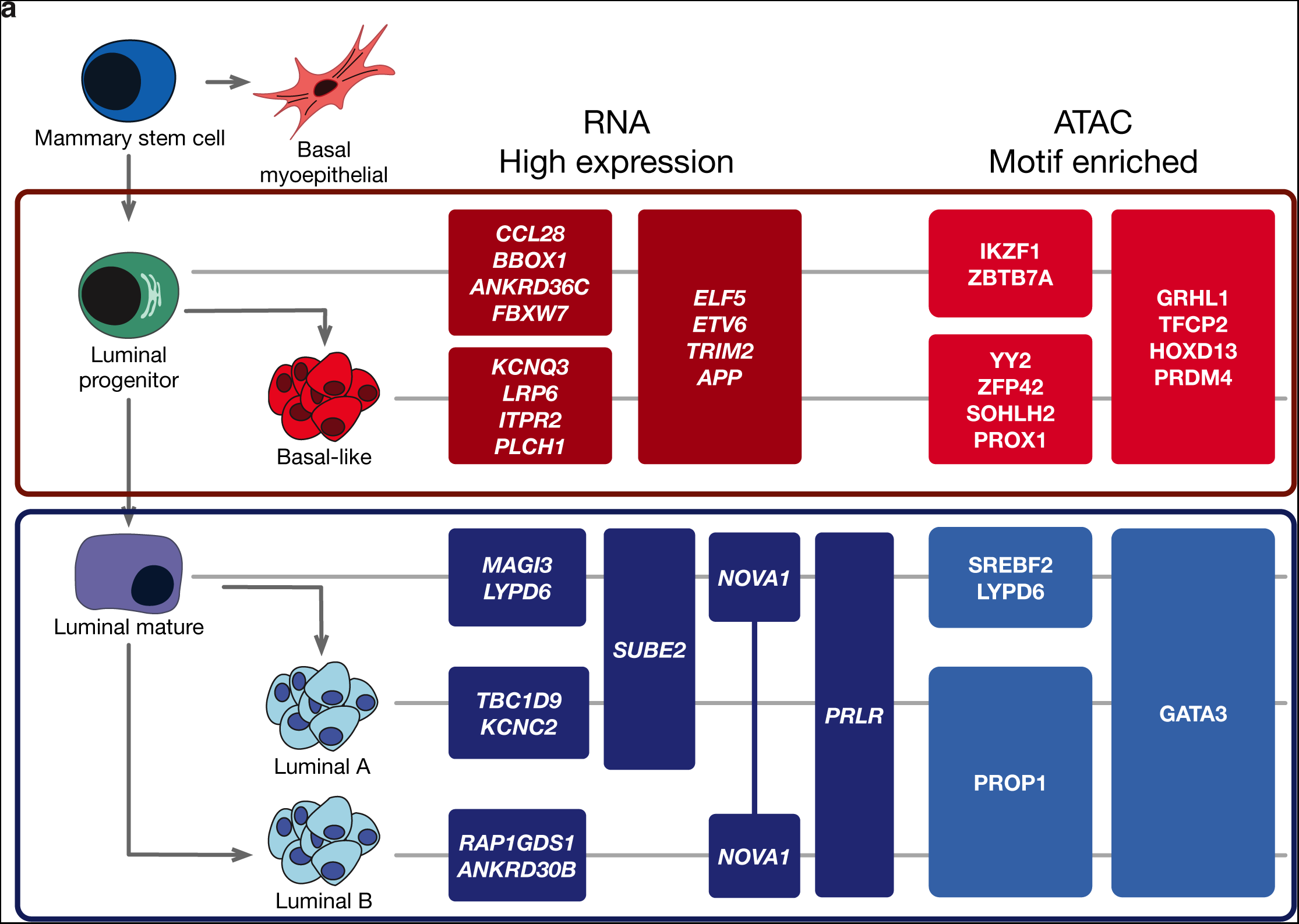
Proposed Model of Breast Cancer Subtype Progression. **a)** Model of proposed cell of origin for subtypes of breast cancer, with key lineage-specific transcription factor motifs and lineage defining expression markers annotated.

## DISCUSSION

Integration of single-cell technologies allows for high-resolution interrogation of tumor subpopulations and stromal and immune components of the tumor microenvironment. Pairing this deep cell-level resolution with multiplex immunofluorescence imaging to provide spatial context, we identified and clarified cell precursors and transitional states and how these transitional populations are associated with chromatin accessibility. Single-nucleus ATAC-seq reveals the transcriptional elements underlying these changes. CODEX immunofluorescence and spatial transcriptomics provide support for these findings and tie single-nucleus findings to discrete histologic structures. Collectively, this study presents multi-omic evidence of the transcriptional programs connecting breast cancer subtypes to distinct cells of origin. Identification of the fine changes associated with transient transitional states is not captured by bulk methods, and may have implications for current treatment paradigms in breast cancer.

Breast cancer subtyping is generally based on bulk gene expression, which is limited by prevalence of non-tumor cell types. In this work, we apply the PAM50 subtyping algorithm in tandem to bulk RNA-seq and snRNA-seq to reliably classify even low-purity tumor samples. Chromatin accessibility from snATAC-seq also clearly separated tumors by subtype and had good concordance with bulk and single nucleus RNA-seq based classifications and highlights transcriptional networks that underlie their gene expression profiles. In addition to known TFs such as GATA3 and FOXP1, we identify HNF1A as a TF specific to luminal A/B breast tumors and luminal mature breast duct cells. HNF1A is not well studied in BC, but is important in colon and pancreatic cancer development, and drives PI3K/AKT signaling in esophageal cancer^72–74^. An antisense product of *HNF1A*, *HNF1A-AS1*, is upregulated in breast cancer and increases proliferation, migration, invasion, and tamoxifen sensitivity in multiple BC cell lines^75^. While this work primarily categorizes breast cancer by PAM50 subtype, there is a lower rate of PR positivity in luminal B compared to luminal A tumors in our data, which could confound findings specific to only one of the luminal subtypes. The lower rate of PR positivity in luminal B tumors is well established ^76^ and further validation could be undertaken in PR positive cohorts. In basal-like BC, we show increased chromatin accessibility for the motifs of known TFs including SOX4, SOX9, and E2F family proliferation-related TFs. Additionally, we highlight TEAD family TF motifs as highly enriched in the open chromatin of basal-like breast cancer, and GRHL1/2 and TFCP2 as enriched in both basal-like tumors and luminal progenitor cells. TEAD TFs are associated with YAP/TAZ transcriptional activators to drive expression of Hippo pathway genes in breast cancer^77^. GRHL2, a member of the Grainhyead TF family, is involved in maintenance of the epithelial phenotype and has been considered a tumor suppressor^78^. However, our results suggest GRHL2 activity is maintained in basal-like tumors. These results are consistent with the proposed oncogenic role of GRHL2: loss of GRHL2 in the BC cell line MCF7 is associated with decreased proliferation, and GRHL2 can also suppress the death receptors FAS and DR5^79,80^. TFCP2 has known oncogenic roles in hepatocellular carcinoma, pancreatic adenocarcinoma, and BC; it has roles in maintaining cell stemness and in EMT and angiogenesis^81^.

Epigenetic features such as DNA methylation and histone modification have been used to track cell lineage and identify cells of origin^28–31^. Because ATAC-seq shows the footprints of TF programs, it complements gene expression data in discerning tumor pathogenesis and carries implications for therapeutic targets to modulate these programs. Chromatin accessibility as assessed by snATAC-seq has key advantages beyond gene expression alone: the maintenance of accessibility patterns across cell types allows for clearer delineation of cell lineage, and it points to upstream effectors of gene expression changes which may themselves be therapeutic targets. In an effort to clarify cells of origin of BC, experimental data robustly support the most widely accepted model that luminal BC arise from a mature ER-positive breast duct cell, whereas basal-like breast cancer arises from a luminal progenitor cell in the same lineage^9,44^. This study adds to our understanding by incorporating many patients with diverse tumor types and mutational spectra and supports the established model that basal-like breast tumors arise from luminal progenitor cells whereas luminal A/B tumors arise from luminal mature cells. The addition of snATAC-seq in many samples adds orthogonal evidence for this conclusion and has not previously been reported. Combining snRNA-seq and snATAC-seq, we implicate several transcriptional programs which are maintained in breast tumors and their proposed cell of origin and have not been extensively reported previously. In both luminal mature duct cells and luminal tumors, BHLHE40 was predicted to regulate co-expressed genes. The precise role of BHLHE40 in BC has not been well described; its role in the luminal lineage may shed light on a targetable pathway in BC treatment or prevention. Further, by evaluating differentially accessible motifs and expression features of each epithelial cell type, we were able to distinguish expression signatures that are altered over the course of tumorigenesis. While we were able to rigorously define the changes between progenitors and tumor cells in luminal A/B and basal-like tumors, we were not able to do so for HER2-enriched tumors. At present we are currently underpowered to address this question likely due to the low sampling size of HER2-enriched tumors in our cohort (3 patients with single-nucleus data). Limited analyses of these few samples shows shared transcriptional features with luminal breast cancer. Future studies focusing on HER2-enriched samples can utilize a similar framework to evaluate the proposed cell of origin for this unique subtype. Additionally, our data include samples from patients receiving a wide variety of systemic therapies as well as treatment-naive samples; additional studies could explore how transcriptional programs in breast cancer are impacted by treatment.

It is well known that substantial immune infiltration is seen in a subset of breast tumors, and that the likelihood of this phenomenon varies by subtype^82,83^. In particular, some basal-like breast tumors are observed to harbor a dense immune infiltrate, and this finding is a positive prognostic feature when it is found^83,84^. Recently, this enrichment in immune infiltration in basal-like BC has led to the approval of pembrolizumab (anti-PD-1 immunotherapy) plus chemotherapy for early-stage TNBC^85^. Using scRNA-seq, we can finely dissect the immune landscape within different breast tumors. We observe significantly more CD8+ exhausted T cells in basal-like tumors compared with other subtypes. Ligand-receptor interaction predicts CTLA4 expressed on CD8+ T cells directly interacts with CD80 or CD86 expressed on multiple myeloid cell types. CCL3 on CD8+ T cells was also predicted to interact with CCR1 or CCR5 on myeloid cells, and CXCL13 on CD8+ T cells was predicted to interact with CXCR5 on B cells or ACKR4 on cancer-associated fibroblasts. All three of these genes, *CTLA4*, *CCL3*, and *CXCL13*, are more highly expressed on CD8+ T cells in basal-like tumors compared to those of other subtypes. CTLA4 provides a negative modulatory signal to T cells interacting with its primary ligands, CD80 or CD86, on an antigen-presenting cell, and is well-described as a key component of tumor immune evasion^86,87^. Anti-CTLA4 blockade with ipilimumab or tremelimumab have been proven effective as immunotherapy in a diverse group of non-breast cancer tumor types, but is not used in standard therapy for breast cancer. The success of the KEYNOTE-522 trial of neoadjuvant pembrolizumab, an anti-PD-1 monoclonal antibody, in TNBC demonstrates a role for immune checkpoint blockade in breast cancer^85^. Our work suggests anti-CTLA4 therapies may also be effective in modulating the anti-tumor immune response, particularly in basal-like breast cancer.

Spatial profiling of cellular proteins with CODEX complements understanding gleaned from single cell or single nucleus-based approaches. In this work, we show GATA3 and cKIT are markers of luminal mature and luminal progenitor cells which are maintained in the transition from benign precursor to invasive breast cancer. Future work will be focused on prospective validation of markers for tumor and benign populations as this may reveal new potential drug targets specific to key populations along the breast cancer evolutionary lineage.

## ACKNOWLEDGEMENTS

We thank the patients, staff, and scientists who contributed to this study as well as the NCI and the HTAN consortium. All HTAN consortium members are named at humantumoratlas.org. We also thank the Siteman Cancer Center and the McDonnell Genome Institute for their support. The following grants supported this work: U2CCA233303 to L.D., R.C.F., W.E.G., and S.T.O.; U24CA211006 to L.D.; U24CA209837 to K.I.S.; R01HG009711 to L.D. and F.C.. Centene Corporation contract (P19-00559) for the Washington University-Centene ARCH Personalized Medicine Initiative to W.E.G.

## AUTHOR CONTRIBUTIONS

**Study Conception & Design**: L.D., W.E.G., R.C.F., F.C., S.A., S.T.O., M.G.C., A.H.K.

**Developed and Performed Experiments or Data Collection**: R.G.J., J.M.H., W.C., S.C., N.N.D., L.T., K.S., A.T.S., S.E.C., M.C., C.B., Y.Z., C.O., M.S., C.C., O.H., A.B., K.L., J.M., F.A., R.A., D.J.V., F.C., A.H., C.S.,

**Computation & Statistical Analysis**: D.C.Z., M.D.I., R.G.J., N.V.T., H.Z., A.K., E.S., C.K.M., H.S., R.L., R.J.L., F.M.R., A.S.S., A.L.N.T.C.,

**Data Interpretation & Biological Analysis**: D.C.Z., M.D.I., R.G.J., N.V.T., A.K., M.A.W., **Writing:** D.C.Z., M.D.I., R.G.J., N.V.T., H.Z., A.K., N.N.D., M.C.W., R.B., S.K.S., A.K., C.M., K.I.S., L.D., M.C., J.M.H., G.A.C., E.S.H., M.G.C., D.J.V.

**Administration**: L.D., W.E.G., R.C.F., R.S.F., C.C.F., L.A.F., S.R.D., E.L.A., T.J., S.A., C.M., K.I.S., J.F.

## FIGURE LEGENDS

**Extended Data Figure 1. InferCNV, nFeature count and epithelial cell type distribution.** a) UMAP plots of copy number events from inferCNV mapped to epithelial cells derived from snRNA data. b) Violin plot of the average nFeature_RNA detected across each sample across three cohorts (one external dataset Wu et al. 2021 and two internal HTAN cohorts). Size of dots indicate the number of cells detected for each sample and boxplot is overlaid on violin plot. c) UMAP representations of epithelial subsets for snATAC and snRNA samples colored by clinical subtype. d) (Left) Barplots indicating proportion of epithelial nuclei per sample identified for the snRNA-seq data. (Right) Barplots indicating proportion of epithelial nuclei per sample identified for the snATAC-seq data.

**Extended Data Figure 2: Exhausted CD8 T cell analysis in snRNA-seq data.** a) UMAP of T cells identified in snRNA-seq data. Cells are colored by cell types. b) Boxplots show the proportion of T cell types relative to all T cells for each piece of tissue separated by subtype and by T cell type. Each point is colored by Treatment Status. Table labeled Wilcoxon test result shows the P value associated with the comparison of proportions of T cells between Group 1 and Group 2. c) Plots showing expression of CCL3, CTLA4, and CXCL13 in Exhausted CD8+ T cells. Size of dot indicates % of cells expressing the gene of interest while color indicates average expression.

**Extended Data Figure 3:** Lymphocyte dense regions in spatial transcriptomics data. Each row indicates a section of a different sample. Left image indicates the lymphocyte dense clusters (L1-LX) selected for evaluating gene expression differences between subtypes. Middle image is the H&E with a region indicated in dashed box that is zoomed in on the right plot to show how we identified lymphocyte dense regions for our analysis.

**Extended Data Figure 4: Spatial mapping of snRNA-seq cell types to Spatial Transcriptomics Data using CytoSPACE.** a) CytoSPACE mapping results of CD4, CD8, Treg and cDC2 to a subset of luminal and basal samples. b) Violin plots of cell type composition of basal enriched cell types. Each grouped violin is separated by cell type and subtype. P-values are derived from stat_compare_means using the method=t.test. c) The heatmap represents the scaled cell type proportion across all breast spatial transcriptomic samples.

**Extended Data Figure 5. TF motifs and pseudotime correlation.** a) Correlation of TF motifs’ scores with pseudotime from precursors to tumor cells from basal-like samples. Color of dot indicates correlation coefficient of each TF separated by sample while the size relates to significance (by FDR). b) Correlation of TF motifs’ scores with pseudotime from precursors to tumor cells from luminal samples.

**Extended Data Figure 6. Histology and snATAC data from MMTV-PyMT model.** a) H&E mouse mammary glands at 12 weeks indicating normal ducts and cancer cells. b) A second H&E of mouse mammary glands at 12 weeks indicating normal ducts and cancer cells. c) UMAP of single-nucleus ATAC-seq data from mouse model. Points are colored by cell type. d) Monocle trajectory analysis of epithelial derived cells from snATAC-seq data. Each point is colored by cell type.

**Extended Data Figure 7. Differential gene expression (DEG) analysis by epithelial lineage.** a) Average expression of differentially expressed genes specific to the basal tumor and/or luminal progenitor cell types. Columns labeled Basal_tumor and Luminal_progenitor indicate whether the gene was identified as a DEG for the respective cell type listed. Heatmap is colored and labeled by average expression of each epithelial cell type for comparison. b) Average expression of differentially expressed genes specific to the Luminal cell types, including: Luminal A tumor, Luminal B tumor, Luminal Mature or Luminal Progenitor. Columns labeled Luminal_progenitor, LumA_tumor, Luminal_mature, LumB_tumor indicate whether the gene was identified as a DEG for the respective cell type listed. Heatmap is colored and labeled by average expression of each epithelial cell type for comparison. c) Dot plot of signac ATAC gene activity values of basal (left) and luminal (right) lineage markers discovered by expression in snRNAseq data. Data is colored by activity value and size of dot is associated with percent of cells with the associated average gene activity score.

**Extended Data Figure 8. Differentially Accessible Motifs (DAM) analysis by epithelial lineage and cell-surface tumor specific markers.** a) Average chromvar motif activity score enriched in the basal tumor and/or luminal progenitor cell types. Columns labeled luminal progenitor (LP), luminal mature (LM), basal myoepithelial (myo), Her_tumor, Lum_tumor and Basal_tumor indicate whether the gene was identified as having a motif score greater than 0 for the respective cell type listed. Heatmap is colored and labeled by the motif activity score of each epithelial cell type for comparison. b) Average chromvar motif activity score enriched in the luminal tumors and/or luminal mature cell types. Columns labeled luminal progenitor (LP), luminal mature (LM), basal myoepithelial (myo), Her_tumor, Lum_tumor and Basal_tumor indicate whether the gene was identified as having a motif score greater than 0 for the respective cell type listed. Heatmap is colored and labeled by the motif activity score of each epithelial cell type for comparison. c) For each gene identified as a tumor specific marker (SYN2, RGS6, SYT1, NPY1R and VTCN1) we have indicated the average expression of the gene listed in each cell type population showing an enrichment in the tumor and progenitor populations relative to other cell types.

**Extended Data Figure 9. Immunofluorescence images of MELK.** a) Immunofluorescence (IF) images of 5 representative regions of HT171B1, b) HT243B1, c) HT271B1, and d) HT308B1. For all images green channel is e-cadherin, blue is DAPI and red is MELK.

**Extended Data Figure 10. Masks associated with Immunofluorescence images of MELK.** a) Masks of the immunofluorescence (IF) images of 5 representative regions of HT171B1, b) HT243B1, c) HT271B1, and d) HT308B1. Masks were generated based on the e-cadherin staining using adaptive thresholding. e) Violin plot of the average pixel intensity of each representative image from the 4 samples. P value indicated comparing the basal samples to luminal samples are derived from stat_compare_means using the method=t.test.

## Supplementary Information

1. **Supplementary Table 1:** Table of clinical, histologic, subtype and cell count metadata
2. **Supplementary Table 2:** Germline variant calls
3. **Supplementary Table 3:** Somatic mutation calls
4. **Supplementary Table 4:** InferCNV calls overlapping whole exome sequencing data
5. **Supplementary Table 5:** Differentially expressed genes in benign epithelial populations
6. **Supplementary Table 6:** Codex Antibody Details

## Methods

### Human Specimens and Clinical Data

All samples were collected with informed consent in concordance with Institutional Review Board (IRB) approval. Primary breast carcinoma samples were collected during surgical resection and verified by standard pathology (IRB protocol 201108117). Blood was collected at the time of surgery into vacuum tubes containing ethylenediaminetetraacetic acid (EDTA) (BD Bioscience). Cells were isolated by ficoll-density centrifugation and frozen in fetal bovine serum with 5% dimethyl sulfoxide. Clinical data was captured in accordance with IRB protocol 20108117, at the time of informed consent and entered into the REDCap database.

### Human Sample Processing

After verification by an attending pathologist, a 1.5 cm x 1.5 cm x 0.5 cm portion of the tumor was removed, photographed, weighed, and measured. Each piece was then subdivided into 6– 9 pieces (depending on the original size) and then further subdivided into four transverse cut pieces. Pieces were then placed into formalin, snap frozen in liquid nitrogen, DMEM, and formalin, respectively.

### Pathologic Parameters and Assessment

Each tumor that is subdivided into smaller increments is subjected to H&E stain and assessed by a pathologist for the following parameters: tumor differentiation and grade, percentage of Tumor Infiltrating Lymphocytes (TIL), lymphovascular invasion and perineural invasion. Tumor viability is also assessed by the presence or absence of necrosis. Both slices of each tumor piece, both L1 and L4 when available were subjected to assessment.

### Mouse Sample Collection and Processing

B6.FVB-Tg(MMTV-PyVT) 634Mul/LellJ (Strain #:022974) female mice were purchased from the Jackson Laboratory. Mice were euthanized by carbon dioxide asphyxiation and five pairs of mouse mammary glands were collected at 12 weeks old. For each pair, the left mammary gland was flash frozen in liquid nitrogen, and the right glands were fixed in 10% neutral buffered formalin (Epredia, 5725) then embedded in paraffin. The 5 left mammary glands were pooled together for snRNA-Seq and snATAC-Seq sample preparation. The paraffin-embedded glands were used for H&E staining. All animal experiments were approved by the Washington University in Saint Louis Institutional Animal Care and Use Committee (IACUC) office.

### Genomic DNA and RNA extraction

Tumor tissues and corresponding normal tissue were obtained from surgically resected specimens and after a piece is removed for fresh single-cell prep the remaining sample is snap-frozen in liquid nitrogen and stored at −80°C. Before bulk RNA/DNA extraction, samples are cryopulverized (Covaris) and aliquoted for bulk extraction methods. Genomic DNA was extracted from tissue samples with either the DNeasy Blood and Tissue Kit (Qiagen, 69504) or the QIAamp DNA Mini Kit (Qiagen, 51304). Total RNA was extracted with TRI reagent (Millipore Sigma, T9424) and treated with DNase I (Qiagen, 79254) using an RNeasy MinElute Cleanup Kit (Qiagen, 74204). RNA integrity was evaluated using either a Bioanalyzer (Agilent Technologies) or TapeStation (Agilent Technologies). Germline genomic DNA was purified from cryopreserved peripheral blood mononuclear cells (PBMCs) using the Qiagen QiaAMP DNA Mini Kit (Qiaggen, 51304) according to the manufacturer’s instructions (Qiagen, Valencia, CA). The DNA quantity was assessed by fluorometry using the Qubit dsDNA HS Assay (Thermo Fisher Scientific, #Q32854) according to manufacturer’s instructions (Thermo Fisher Scientific, Waltham, MA)

### Whole Exome Sequencing

100-250ng of genomic DNA was fragmented on the Covaris LE220 instrument targeting 250bp inserts. Automated dual indexed libraries were constructed with the KAPA Hyper library prep kit (Roche) on the SciClone NGS platform (Perkin Elmer). Up to ten libraries were pooled at an equimolar ratio by mass prior to the hybrid capture targeting a 5µg library pool. The library pools were hybridized with the xGen Exome Research Panel v1.0 reagent (IDT Technologies) that spans 39 Mb target region (19,396 genes) of the human genome. The libraries were hybridized for 16-18 hours at 65°C followed by stringent wash to remove spuriously hybridized library fragments. Enriched library fragments were eluted and PCR cycle optimization was performed to prevent over amplification. The enriched libraries were amplified with KAPA HiFi master mix (Roche) prior to sequencing. The concentration of each captured library pool was accurately determined through qPCR utilizing the KAPA library Quantification Kit according to the manufacturer’s protocol (Roche) to produce cluster counts appropriate for the Illumina NovaSeq-6000 instrument. 2x150 paired end reads were generated targeting 12Gb of sequence to achieve ∼100x coverage per library.

### RNA Sequencing

Total RNA integrity was determined using Agilent Bioanalyzer or 4200 Tapestation. Library preparation was performed with 500ng to 1ug of total RNA. Ribosomal RNA was blocked using FastSelect reagents (Qiagen) during cDNA synthesis. RNA was fragmented in reverse transcriptase buffer with FastSelect reagent and heating to 94 degrees for 5 minutes, 75 degrees for 2 minutes, 70 degrees for 2 minutes, 65 degrees for 2 minutes, 60 degrees for 2 minutes, 55 degrees for 2 minutes, 37 degrees for 5 minutes, 25 degrees for 5 minutes. mRNA was reverse transcribed to yield cDNA using SuperScript III RT enzyme (Life Technologies, per manufacturer’s instructions) and random hexamers. A second strand reaction was performed to yield ds-cDNA. cDNA was blunt ended, had an A base added to the 3’ ends, and then had Illumina sequencing adapters ligated to the ends. Ligated fragments were then amplified for 15 cycles using primers incorporating unique dual index tags. Fragments were sequenced on an Illumina NovaSeq-6000 S4 instrument generating approximately 30M paired end 2x150 reads per library.

### Single-cell Suspension Preparation

For each tumor approximately 15-100 mg of 2-4 sections of each tumor and/or normal piece of tissue were cut into small pieces using a blade and processed separately. Enzymes and reagents from the human tumor dissociation kit (Miltenyi Biotec: 130-095-929) were added to the tumor tissue along with 1.75 mL of DMEM. Resulting suspension was loaded into a gentleMACS C-tube (Miltenyi Biotec: 130-093-237) and subject to the gentleMACS Octo Dissociator with Heaters (Miltenyi BiotecL 130-096-427). After 30 to 60 minutes on the heated dissociation program (37h_TDK_1), samples are removed from the dissociator and filtered through a 40 uM Mini-Strainer (PluriSelect # 43-10040-60) or Nylon mesh 40um (Fischer Scientific: 22-363-547) into a 15 mL conical tube on ice. The sample is then spun down at 400 g for 5 min at 4℃. After removing supernatant, when a red pellet was visible the cell pellet was resuspended using 200 uL to 3 mL of ACK Lysis Solution (ThermoFisher: A1049201) for 1 to 5 minutes. To quench the reaction, 10 mL of PBS (Corning, 21-040-CM) with 0.5% BSA (Miltenyi Biotec: 130-091-376) is added and spun down at 400 g for 5 min at 4℃. After removing supernatant, cells are resuspended in 1 mL of PBS (Corning: 21-040-CM) with 0.5% BSA, live and dead cells are visualized using Trypan Blue. If greater than 40% of dead cells are present, the sample is spun down at 400 g for 5 min at 4℃ and subject to the dead cell removal kit (Miltenyi Biotec, 130-090-101). Finally the sample is spun down at 400 g for 5 min at 4℃ and resuspended in 500 μL to 1 mL of PBS with 0.5% BSA to a final concentration of 700 to 1,500 cells per μl.

### Single-cell library prep and sequencing

Utilizing the Chromium Next GEM Single Cell 3’ GEM, Library & Gel Bead Kit v3.3 and Chromium instrument, approximately 17,500 to 25,000 cells were partitioned into nanoliter droplets to achieve single cell resolution for a maximum of 10,000-15,000 individual cells per sample (10X Genomics). The resulting cDNA was tagged with a common 16nt cell barcode and 10nt Unique Molecular Identifier during the RT reaction. Full length cDNA from poly-A mRNA transcripts was enzymatically fragmented and size selected to optimize the cDNA amplicon size (approximately 400 bp) for library construction (10x Genomics). The concentration of the 10x single cell library was accurately determined through qPCR (Kapa Biosystems) to produce cluster counts appropriate for the HiSeq 4000 or NovaSeq 6000 platform (Illumina). 26x98bp sequence data were generated targeting 50K read pairs/cell, which provided digital gene expression profiles for each individual cell.

### Single-nuclei RNA and ATAC library preparation and sequencing

About 20-30 mg of cryopulverized powder from BRCA specimens was resuspended in 2ml Lysis buffer (10 mM Tris-HCl (Thermofisher: 15567027) (pH 7.4); 10 mM NaCl (Themofisher: AM9759); 3 mM MgCl2 (Thermofisher: AM9530G); and 0.1% NP-40 (Sigma: 74385-1L)) plus 0.1 U/ul RNase Inhibitor (Invitrogen: AM2696). This suspension was pipetted gently for 6-8 times, incubated on ice for 30 seconds, and pipetted again for 4-6 times. The lysate containing free nuclei was filtered through a 40 μm cell strainer. We washed the filter with 1 mL Wash and Resuspension buffer (1X PBS (Corning: 21-040-CM) + 2% BSA (Miltenyi Biotec: 130-091-376) + 0.2 U/μL RNase inhibitor) plus 0.1 U/ul RNase Inhibitor and combined the flow through with the original filtrate. After a 6-minute centrifugation at 500 x g and 4 °C, the nuclei pellet was resuspended in 300 μL of Wash and Resuspension buffer plus 0.1 U/ul RNase Inhibitor. After staining with 1 μL 7AAD (ATAC or multiome) or DRAQ5 (RNA) the nuclei were further purified by Fluorescence Activated Cell Sorting (FACS). FACS-purified nuclei were centrifuged again and resuspended in a small volume (about 30 μL of Wash and Resuspension buffer plus 0.1 U/ul RNase Inhibitor). After counting and microscopic inspection of nuclei quality, the nuclei preparation was diluted to about 1,000 nuclei/μL. About 20,000 nuclei were used for single-nuclei RNA sequencing (snRNA-seq) by the 10X Chromium platform. For snRNA, we loaded the single nuclei onto a Chromium Next Gem Chip G Kit and processed them through the Chromium Controller to generate GEMs (Gel Beads in Emulsion). For a subset of samples with joint snRNA and snATAC the multiome kit, Chromium Next GEM Single Cell Multiome ATAC + Gene Expression was used. For ATAC only samples FACS-purified nuclei were centrifuged again and resuspended in 5 μL of Diluted Nuclei Buffer. After counting and microscopic inspection of nuclei quality, the nuclei preparation, about 10,000 nuclei were used for single-nuclei ATAC sequencing (snATAC-seq) by the 10X Chromium platform. We loaded the single nuclei onto a Chromium Next Gem Chip H Kit and processed them through the Chromium Controller to generate GEMs (Gel Beads in Emulsion). After that, post GEM-RT Cleanup was performed with target cell recovery ≥2,000. We then prepared the sequencing libraries following the manufacturer’s protocol. All sequencing was performed on an Illumina NovaSeq 6000 S4 flow cell. The libraries were pooled and sequenced using the XP workflow according to the manufacturer’s protocol with a 28x8x98bp sequencing recipe. The resulting sequencing files were available as FASTQs per sample after demultiplexing.

### Fluorescence-activated cell sorting (FACS)

Depending on the pellet size, 100-500 uL of nuclei suspension in the wash buffer (2% BSA + 1x PBS + RNase inhibitor) was stained with DRAQ5 or 7AAD for RNA or ATAC sequencing, respectively (7AAD was used for multiome processing). Namely, snRNA-seq nuclei were stained with 1 uL of DRAQ5 per 300 uL of the sample, and snATAC-seq nuclei were stained with 1uL of 7AAD per 500 uL of the sample. Sorting gates were based on size, granularity, and dye staining signal.

### Immunofluorescence and microscopy

Fresh tissues were fixed in 10% neutral buffered formalin (Epredia, 5725) at room temperature overnight but less than 24 hours. Tissues were then dehydrated, infiltrated with wax, and embedded into paraffin blocks. After tissues were processed into Formalin-fixed paraffin-embedded (FFPE) blocks, 5 μm sections were cut and placed on glass slides. Next, sections were deparaffinized and rehydrated, followed by antigen retrieval using Tris EDTA buffer pH 9 (Genemed, 10-0046) or 1x sodium citrate at pH 6 (Sigma, C9999) according to manufacturer’s recommendation for specific antibodies. Then, sections were blocked with 100 mM glycine for 20 minutes, followed by blocking with 10% normal serum and 1% BSA for 1 hour at room temperature. A negative control and a secondary antibody control were used in each experimental setting. Primary antibodies for MELK (Thermofisher, catalog: MA517120) and Ecadherin (R&D, catalog: AF748) were applied on sections at 4°C overnight, followed by the incubation of appropriate secondary antibodies the next day. Images were collected by a Leica DMi8 microscope.

### CODEX preparation and imaging

Carrier-free monoclonal or polyclonal anti-human antibodies were purchased from different companies (Supplementary Table 6) and verified using immunofluorescence (IF) staining in multiple channels. After the screening, antibodies were conjugated using Akoya Antibody Conjugation Kit (Akoya Biosciences, SKU 7000009) with a barcode (Akoya Biosciences) assigned based on the IF staining results. Several common markers were directly purchased through Akoya Biosciences (Supplementary Table 6. CODEX staining and imaging were performed according to the manufacturer’s instruction (CODEX User Manual -Rev C). Briefly, 5 μm FFPE sections were placed on APTES (Sigma, #440140) coated coverslips and baked at 60°C overnight before deparaffinization. The next day, tissues were incubated in in xylene, rehydrated in ethanol, and washed in ddH2O before antigen retrieval with TE buffer, pH 9 (Genemed, #10-0046) in the boiling water for 10 min in a rice cooker. The tissues were then blocked using the blocking buffer (CODEX staining kit, SKU 7000008) and stained with the marker antibody panel (Supplementary Table 6) to a volume of 200 µl for 3 hours at room temperature in a humiliated chamber. Imaging of the CODEX multicycle experiment was performed using Keyence fluorescence microscope (model BZ-X810) equipped with a Nikon CFI Plan Apo λ 20x/0.75 objective, the CODEX instrument (Akoya Biosciences, USA), and CODEX Instrument Manager (CIM) (Akoya Biosciences, USA). Exposure times, dilutions and the order of markers per cycle are listed in Supplementary Table 6. The raw images were then stitched and processed using the CODEX processor (Akoya Biosciences, USA). After multiplex imaging was completed, hematoxylin and eosin (H&E) staining was performed on the same tissue.

### Spatial Transcriptomics prep and sequencing

OCT-embedded tissues were cryosectioned and placed on Visium Spatial Gene Expression Slide following Visium Spatial Protocols-Tissue Preparation Guide (10X Genomics, CG000240 Rev A). Briefly, fresh tissues were coated carefully and thoroughly with room temperature OCT without any bubbles. OCT-coated tissues were then placed on a metal block chilled in dry ice until the OCT turned solidified and white. After RNA quality check using Tapestation and morphology check using H&E staining for the OCT-embedded tissues, blocks were scored into proper size that fit the Capture Areas and then sectioned into 10 μm sections. After the tissue placement into the Capture Area, sections were fixed in methanol, stained with hematoxylin and eosin, and imaged at 20x magnification using the brightfield imaging setting on Leica DMi8 microscope. Tissues were then permeabilized for 18 minutes and Spatial Transcriptomics libraries were constructed following Visium Spatial Gene Expression Reagent Kits User Guide CG000239 Rev A (10X Genomics). Briefly, cDNA was reverse transcribed from the poly-adenylated mRNA which was captured by the primers on the slides. Next, the second strand was synthesized and denatured from the first strand. Free cDNA were then transferred from slides to tubes for further amplification and library construction. Libraries were sequenced on the S4 flow cell of Illumina NovaSeq 6000 system.

## QUANTIFICATION AND STATISTICAL ANALYSIS

### Genomic Data Analysis

#### Tumor-Normal Somatic Variant Calling

Somatic variants were called from WES tumor and normal paired BAMs using somaticwrapper v1.6, a pipeline designed for detection of somatic variants from tumor and normal exome data. The pipeline merges and filters variant calls from four callers: Strelka v2.9.2^88^, VarScan v2.3.8^89^, Pindel v0.2.5^90^, and MuTect v1.1.7^91^. SNV calls were obtained from Strelka, Varscan, and Mutect. Indel calls were obtained from Stralka2, Varscan, and Pindel. The following filters were applied to get variant calls of high confidence: normal VAF ≤ 0.02 and tumor VAF ≥ 0.05, read depth in tumor ≥ 14 and normal ≥ 8, indel length < 100 bp, all variants must be called by 2 or more callers, all variants must be exonic, and exclude variants in dbSNP but not in COSMIC.

#### Tumor-Only Somatic Variant Calling

Tumor-only somatic variants were called using Mutect2 (v4.1.2.0) best-practice pipeline (gatk.broadinstitute.org) with the GDC Panel of Normal (PON) data (gdc.cancer.gov/about-data/data-harmonization-and-generation/gdc-reference-files; gatk4_mutect2_4136_pon.vcf.tar). To further reduce false positives, we kept the mutation sites with ≥ 20X coverage, > 3 reads, and ≥ 0.1 tumor VAF, which was supported by bam-readcount (https://github.com/genome/bam-readcount).

#### Somatic Variant Rescue

In some tumor cases, we called somatic variants of driver genes for some pieces, but not for all pieces. Therefore, we used bam-readcount to rescue those variants for the piece (s) without them. We kept default parameters for bam-readcount, except setting --min-mapping-quality as 20 and --min-base-quality as 20.

#### Germline Variant Calling and Annotation

Germline variant calling was performed using an in-house pipeline germlinewrapper v1.1, which implements multiple tools for the detection of germline INDELs and SNVs. Germline SNVs were identified using VarScan v2.3.8 (with parameters: --min-var-freq 0.10 --p-value 0.10, --min-coverage 3 --strand-filter 1) operating on a mpileup stream produced by samtools v1.2 (with parameters: -q 1 -Q 13) and GATK v4.0.0.0 ^92^ using its haplotype caller in single-sample mode with duplicate and unmapped reads removed and retaining calls with a minimum quality threshold of 10. Germline INDELs were identified using VarScan (version and parameters as above) and GATK (version and parameters as above) in single-sample mode. Single nucleotide variants (SNVs) were based on the union of raw GATK and VarScan calls. We required that indels were called by Pindel or at least two out of the three callers (GATK, VarScan, Pindel). Cutoffs of minimal 10X coverage and 20% VAF were used in the final step to report the high-quality germline variants. All resulting variants were limited to the coding region of the full-length transcripts obtained from Ensembl release 100 plus additional two base pairs flanking each exon to cover splice donor/acceptor sites. We also required variants to have allelic depth ≥ 5 for the alternative allele in both tumor and normal samples and filtered out any indels longer than 100bp.

#### Germline Variant Pathogenic Classification

Germline variants called with GermlineWrapper were annotated with the Ensembl Variant Effect Predictor (VEP)^93^(version 100 with default parameters, except where --everything) and their pathogenicity was determined with our automatic pipeline CharGer^49^ (version 0.5.4 with default CharGer scores, https://github.com/ding-lab/CharGer/tree/v0.5.4), which annotates and prioritizes variants based on the American College of Medical Genetics and Genomics - Association for Molecular Pathology (ACMG-AMP) guidelines^94^. The detailed implementation, score of each evidence level, and parameters used are as previously described^95^ .

We selected rare variants with ≤ 0.05% allele frequency (AF) in gnomAD (r2.1.1) or 1000 Genomes^96^. We also performed read count analysis using bam-readcount (https://github.com/genome/bam-readcount; version 0.8 with parameters -q 10, -b 15) in both normal and tumor samples. We required variants to have at least 5 counts of the alternative allele and a variant allele frequency (VAF) of at least 20% in both tumor and normal. Variants affecting known cancer predisposition genes (previously described in^95^) were manually reviewed with the Integrative Genomics Viewer (IGV) software (21221095; version 2.8.2). We considered variants to be pathogenic (P) if they were known pathogenic variants in ClinVar; likely pathogenic (LP) if CharGer score > 8; and prioritized variant of uncertain significance (PVUS) if CharGer score > 4.

Variants called in cases where a normal sample was not available (tumor only) were further filtered for removal of potential somatic events. Variants in these cases called by our germline pipeline which were also called by our somatic pipeline were filtered out, as well as variants not previously reported in gnomAD or reported in COSMIC (extracted from VEP annotation).

#### RNA Quantification

We used an in-house bulk RNA Seq analysis pipeline for quantification (https://github.com/ding-lab/HTAN_bulkRNA_expression). Briefly, for each sample, the raw sequence reads are aligned into BAM files using STAR(2.7.4a) 2-pass alignment with GRCh38 as the reference. The resulting BAM files are then quantified as a raw count-matrix using feature counts (subread, version 2.0.1). For both alignment and quantification, gene annotations are based on Gencode v34. The raw counts are then converted to FPKM-UQ based on GDC’s formula and then log2 transformed with 1 pseudocount (https://docs.gdc.cancer.gov/Data/Bioinformatics_Pipelines/Expression_mRNA_Pipeline/#upper-quartile-fpkm).

#### Expression-based Subtyping

Bulk expression subtyping is performed according to methods detailed in^97^ using the log2 upper quartile normalized FPKM reads for all bulk RNA-seq samples. Median values of all 50 PAM50 genes were provided for median adjustment. To minimize the influence of any one sample on subtype calls, median values for the 50 PAM50 genes were bootstrapped using a subset of the bulk RNA-seq data comprising all 17 ER-negative samples and an equal number of randomly-selected ER-positive samples. PAM50 subtype assignments were called for all samples using 1,000 such sets of median values, and a final subtype assignment for each sample was the subtype most commonly called across these iterations. The PAM50 subtype-calling algorithm was run using code provided in^97^ using R version 4.0.2.

### scRNA-seq and snRNA-seq Quantification and Analysis

#### scRNA-seq Data Preprocessing

For each sample, we obtained the unfiltered feature-barcode matrix per sample by passing the demultiplexed FASTQs to Cell Ranger v3.1.0 ‘count’ command using default parameters and the prebuilt GRCh38 genome reference 3.0.0 (GRCh38 and Ensembl 93).

Seurat v4.1.0^98,99^ was used for all subsequent analysis. First, a series of quality filters were applied to the data to remove those barcodes which fell into any one of these categories recommended by Seurat: too few total transcript counts (< 300); possible debris with too few genes expressed (< 200) and too few UMIs (< 1,000); possible more than one cell with too many genes expressed (> 10,000) and too many UMIs (> 10,000); possible dead cell or a sign of cellular stress and apoptosis with too high proportion of mitochondrial gene expression over the total transcript counts (> 10%). Doublets were filtered out using Scrublet (https://github.com/AllonKleinLab/scrublet). Scrublet was run on each sample separately with the following parameter settings: expected_doublet_rate=0.06, min_counts=2, min_cells=3, min_gene_variability_pctl=85, n_prin_comps=30. The doublet score threshold was adjusted manually, which can separate the two peaks of a bimodal simulated doublet score histogram. We constructed a Seurat object using the unfiltered feature-barcode matrix for each sample. Each sample was scaled and normalized using Seurat’s ‘SCTransform’ function to correct for batch effects (with parameters: vars.to.regress = c(“nCount_RNA”, “percent.mito”), variable.features n = 2000). Any merged analysis or subsequent subsetting of cells/samples underwent the same scaling and normalization method. Cells were clustered using the original Louvain algorithm and top 30 PCA dimensions via ‘FindNeighbors’ and ‘FindClusters’ (with parameters: resolution = 0.5) functions. The resulting merged and normalized matrix was used for the subsequent analysis.

#### scRNA-seq and snRNA-seq Cell Type Annotation

Main cell types were assigned to each cluster by manually reviewing the expression of a comprehensive set of marker genes. These assignments were all done by one person to maximize consistency. The marker genes used were *MYC, ESR1, ESR2, CD24, AR, PGR, PECAM1, EZH2, EGFR, TP53, PARP1, FAP, BCL2, RB1, CA9, CASP3, CDH1, AKT1, ERBB2, EPCAM, BIRC5, MET, IGHG1, PTEN, CD44, VIM, KRT14, CD74, MKI67, PTPRC, SMN1, SMN2, ITGA6, KRT1, CCNB1, CD3G, CD68, KRT8, KRT18, KRT19, KRT5, KRT7* (tumor); *KIT, SLPI, ALDH1A3, CD14* (luminal progenitor); *AGR2*, *STC2*, *ANRKD30A* (luminal mature); *OXTR, ACTA2*, *KRT14*, *KRT17*, *FST*, *PDPN*, *COL7A1* (basal/myoepithelial); *VWF, PECAM1, FLT4, FLT1, FLT3, KDR, PLVAP, ANGPT2, TRIM24, ACTA2* (endothelial); *PTPRG, FLT1, VEGFA, NRP1, NKAIN2, PECAM1* (vascular_endothelial_cells); *EPCAM, AMBP, MUC1* (epithelial); *TIMP1, FN1, POSTN, ACTA2, BST2, LY6D, COL6A1, SLC20A1, COL6A2, KRT16, CD9, S100A4, EMP1, LRRC8A, EPCAM, PDPN, ITGB1, PDGFRA, THY1* (fibroblast); *ESAM, ATP1B2, ANO1, GM13889, DES, AOC3, NDUFA4L2, GUCY1A3, GDPD3, MCAM, HIGD1B, CPE, KCNJ8, ABCC9, RGS4, SPARCL1, RGS5* (vCAF); *SMOC2, ITGBL1, FBLN1, CDH11, CRABP1, PDGFRA, SVEP1, PDPN, LSP1, CPXM1, LRRC15, CILP, DCM, LUM, MFAP5, FBLN2, OLFML3, RNASE4* (mCAF); *ANIN, CDCA3, TPX2, CDCA8, FAM64A, NUF2, BIRC5, CEP55, SKA1, KIF15, TTK, MELK, TOP2A, PBK, CCNA2, SPC25, MKI67* (cCAF); *SDC1, IGHG1, IGHG3, IGHG4, IGHA1, IGHA2, FCRL5, IGKC, TNFRSF17, TNFRSF13B, IGLC2* (plasma); *CD19, MS4A1, CD79A, CD79B, CD83, CD86 (B cells); CD8B, CD8A, CD3E, CD3D* (CD8 T cells); *CD4, CD3E, CD3D, most show SELL, CCR7, IL7R, TCF7 and LEF1 expression* (CD4 T cells); *XCL2, XCL1, SPON2, KLRF1, KIR2DL3, IL2RB, HOPX, CLIC3, CD7, KLRB1, KLRD1, GZMA, PRF1, CD160, NCAM1, FCGR3A* (NK); *NK markers and CD3E, CD3D* (NKT cells); *FCER1A, KIT, FCER2, ENPP3* (mast); *PNPLA2, CAV1, FABP4, COX4I1, LMNB1, PPARG, CEBPA, LEP, HOXC8, HOXC9, UCP1, CIDEA, PRDM16, ZIC1, LHX8, MPZL2, EPSTI1, TNFRSF9, TMEM26, TBX1, CITED1, SHOX2, SLC7A10, SLC36A2, P2RX5* (adipocytes); *ITGAM, LGALS3, CD68, CD163, LYZ, ADGRE1, LAMP2* (macrophages); *CD14, FCGR3A, FCGR1A* (monocytes); *IL3RA, HLA-DRA, CLEC4C, NRP1, PTPRC, LILRA4, TLR7, TLR9, IRF7, GZMB* (pDC); *CD1C, CD83, THBD, CD209, CD8A, CD8B, ITGAM, ITGAX, ITGAE, LY75, CD74, CD86, HLA-DPA1* (cDC); *CD1C, CD1A, FLT3, ZBTB46* (cDC2)*; XCR1, CLEC9A, IRF8, FLT3, ZBTB46, BATF3* (cDC1); We further subdivided certain cell types into subtypes or cell states using the following: *IKZF2, TNFRSF18, FOXP3, CTLA4, IL7R, IL2RA* (Treg); *GZMH, GZMB, GZMA, PRF1, IFNG, FASLG, LAMP1, CD8A, CD8B, CD3E, CD3D* (cytotoxic T cell); *GZMK* (pre-exhausted T-cell); *VSIR, TIGIT, ICOS, EOMES, HAVCR2, PDCD1, BTLA, CD244, LAG3, CD160, CTLA4, CD96* (exhausted T cell); *CD69*, *CD28*, *CD44*, *DPP4* (activated T cell); high ribosomal gene expression (RPhigh CD4 T cell); *IL1A*, *IL1B* (IL1A+ macrophages); high *TLR2* and *CD86* and low *CD163* and *MRC1* (M1 macrophages); high *CD163*, high *MRC1* (M2 macrophages); high *CD163*, *MRC1*, *TLR2*, and *CD86* (M1/M2 macrophages); high *CD163* and low *MRC1* (M2 partial macrophages); *MKI67* (proliferation marker); high *CD69*, medium-low *CCR7*, medium-low *SELL*, medium-low *IL7R* (activated T-cell).

#### snRNA-seq Mouse Cell Type Annotation

To annotate the mouse single-cell data the following markers were used: Basal = Epcam, Krt5, Acta2, Myh11, Krt14, Trp63, Krt17, Myl9; Luminal Mature = Areg, Cited1, Ly6d, Prlr, Esr1, Pgr, S100a6; Luminal Progenitor = Kit, Aldh1a3, Cd14, Gabrp, Tspan8; Cycling_cells = Birc5, Hmgb2, Stmn1, Mki67; Immune = Ptprc, Fyb; Stroma = Col4a1, Sparc, Col4a2, Lamb1, Col5a2; Fibroblast = Fabp4, Lpl, C4b, Mylk, Hk2, Slc4a4, Dio2, Vegfa.

#### snRNA-seq PAM50 Subtype Assignment

For single-nuclei data, to overcome data sparseness, expression-based subtyping was performed at the cluster level. After cell type annotation of the combined Seurat object across all samples, a separate Seurat object was created comprising only tumor cells and was clustered again using the original Louvain algorithm and top 30 PCA dimensions via ‘FindNeighbors’ and ‘FindClusters’ (with parameters: resolution = 0.5) functions. Mean expression of the 50 genes used in the PAM50 algorithm were obtained per cluster. PAM50 subtype assignments were obtained for each of these cluster means in the same way as was done for bulk RNA-seq data (without bootstrapping).

#### scRNA-seq and snRNA-seq Data Preprocessing

For each sample, we obtained the unfiltered feature-barcode matrix per sample by passing the demultiplexed FASTQs to Cell Ranger v3.1.0 ‘count’ command using default parameters and a customized pre-mRNA GRCh38 genome reference to capture both exonic and intronic reads. The customized genome reference modified the transcript annotation from the 10x Genomics pre-built human genome reference 3.0.0 (GRCh38 and Ensembl 93). Seurat v4.1.0 ^98,99^ was used for all subsequent analysis. First, a series of quality filters were applied to the data to remove those barcodes which fell into any one of these categories recommended by Seurat: too few total transcript counts (<300); possible debris with too few genes expressed (<200) and too few UMIs (<1,000); possible more than one cell with too many genes expressed (>10,000) and too many UMIs (>10,000); possible dead cell or a sign of cellular stress and apoptosis with too high proportion of mitochondrial gene expression over the total transcript counts (>10%).We constructed a Seurat object using the unfiltered feature-barcode matrix for each sample. Each sample was scaled and normalized using Seurat’s ‘SCTransform’ function to correct for batch effects (with parameters: vars.to.regress = c(“nCount_RNA”, “percent.mito”), variable.features.n = 3000). We then merged all samples and repeated the same scaling and normalization method. All cells in the merged Seurat object were then clustered using the original Louvain algorithm (Blondel et al., 2008) and top 30 PCA dimensions via ‘FindNeighbors’ and ‘FindClusters’ (with parameters: resolution = 0.5) functions. The resulting merged and normalized matrix was used for the subsequent analysis.

#### Gene regulatory networks

To infer gene regulatory networks of transcription factors, we used pySCENIC interface version (v0.11.2) from the SCENIC pipeline^100^. We applied SCENIC on SCT-normalized assay of sampled merged snRNA combo object, 500 cells sampled randomly per cell type of each sample. The first step of SCENIC workflow is utilizing a regression per-target approach, GRNBoost2, to infer coexpression modules. We provided the list of unique TFs that are present in the JASPAR2020 db^101^ as input. Steps 2 and 3 of regulon prediction were run with default parameters and using RcisTarget hg38 refseq_r80 v.9 gene-motif ranking databases (10 kbp around the TSS, and 500 bp around TSS). Next, we re-calculated the AUCell score, the regulon activity, to identify significant regulons (TFs) in each cell type. Then only regulons (Transcription factors) with at least 20 regulated genes were considered in the final heatmap. Finally, we generated a binary regulon activity heatmap to show gene regulatory networks relationships between Transcription factors and their target genes by using ComplexHeatmap R package.

#### Mapping and quantification of snATAC-seq and snMultiome-seq

To process sequenced snATAC-seq and snMutiome-seq data, we used cellranger-atac count (v.2.0, 10X Genomics) and cellranger-arc count (v.2.0, 10X Genomics) pipelines respectively. These pipelines allow to filter and map snATAC-seq reads and to identify transposase cut sites, and cellranger-arc count pipeline also performs filtering and alignment of snRNA-seq reads. The GRCh38 human reference was used for the reads mapping.

#### Peak calling for snATAC-seq data

We performed peak calling using MACS2^102^. We further removed peaks from the Y-chromosome, and also the ones that overlap genomic regions containing “N”. All peaks were resized to 501 bp centered at the peak summit defined by MACS2. After this, we performed an iterative removal procedure described in Corces et al.^103^ in order to get the set of non-overlapping peaks. In brief, we start with retaining the most significant peak by MACS2 peak score (−log10(q-value)) and remove all peaks that have direct overlap with it. We repeat this procedure for the remaining peaks, until we have the set of non-overlapping peaks (sample peak set). The resulting set of sample peaks was used to calculate peak-count matrix using FeatureMatrix from the Signac package v.1.3.0 (https://github.com/timoast/signac), which was also used for downstream analysis.

#### QC of snATAC-seq data

QC-filtering of the snATAC-datasets was performed using functions from the Signac package. Filters that were applied for the cell calling include: number of fragments in peaks > 1,000 and < 20,000, percentage of reads in peaks > 15, ENCODE blacklist regions percentage <0.05 (https://www.encodeproject.org/annotations/ENCSR636HFF/), nucleosome banding pattern score < 10, enrichment-score for Tn5-integration events at transcriptional start sites > 2. Peaks were annotated using R package ChiPseeker ^104^ using transcript database TxDb.Hsapiens.UCSC.hg38.knownGene. The promoter region was specified (−1000,100) relative to the transcription start site.

#### Normalization, feature selection, and dimension reduction of snATAC-seq data

The filtered peak-count matrix was normalized using term frequency-inverse document frequency (TF-IDF) normalization implemented in the Signac package. This procedure normalizes across cells, accounting for differences in coverage across them and across peaks and giving higher values to the more rare peaks. All the peaks were used as features for the dimensional reduction. We used the RunSVD Signac function to perform singular value decomposition on the normalized TF-IDF matrix, which is known as Latent Semantic Indexing (LSI) dimension reduction. The resulting 2:30 LSI components were used for non-linear dimension reduction using function RunUMAP from the Seurat package.

#### Clustering of snATAC-seq data

The nuclei were clustered using a graph-based clustering approach implemented in Seurat. First, we utilized the Seurat function FindNeighbors to construct a Shared Nearest Neighbor graph using the 2:30 LSI components. Next, we used the FindClusters function to iteratively group nuclei together while optimizing modularity using the Louvain algorithm.

#### Merging of snATAC-seq data across samples

Merging of snATAC-seq datasets was performed using functions from the Signac and Seurat packages. In order to normalize peaks’ significance scores across samples and cancers, we converted MACS2 peak scores (−log10(q-value)) to a score per million as was described in Corces et al ^103^. To get the set of peaks for merging, we first combined peaks from all samples of the cohort. For overlapping peaks across samples, we performed an iterative removal procedure, the same as was used for creating individual sample sets of peaks, using normalized peak scores as described above. By this procedure we obtained the cohort level peak set. The resulting list of peaks was quantified and was used to create a peak-cell matrix so that the set of features was the same across all snATAC-seq samples. After that, the merge function from the Seurat package was used to merge snATAC-seq datasets. Next, we performed TF-IDF normalization. LSI-dimensional reduction was performed using the RunSVD function. Non-linear dimension reduction was performed using the RunUMAP function with the first 2:50 LSI-components.

#### Cell type label transfer from snRNA-seq to snATAC-seq data

Cell type label transfer was performed using functions from Signac and Seurat. First, we quantified chromatin accessibility associated with each gene by summing the reads overlapping the gene body and its upstream region of 2Kb, thus creating the gene by cell matrix. Coordinates for the genes were used from the Ensembl database v.86 (EnsDb.Hsapiens.v86 package). Next, we performed log-normalization of the resulting matrices using the NormaliseData function. The integration of paired snATAC-seq and snRNA-seq datasets was performed using the FindTransferAnchors function with the Canonical Correlation Analysis (CCA) option for the dimensional reduction. We then utilized the TransferData function to transfer cell type labels from the snRNA-seq dataset to the snATAC-seq dataset using the obtained set of anchors from the previous step. Then cell types were re-evaluated at the cohort object level, where for each cluster cell type label was assigned by the most abundant cell type in that cluster.

#### Identifying differentially accessible chromatin regions using snATAC-seq data

To identify cancer specific differentially accessible chromatin regions (DACRs), we performed a comparison for tumor cells from each cancer vs tumor cells from all other cancers. For this we used FindMarkers function from the Seurat package with logistic regression test and the fraction of fragments peaks as a latent variable to reduce the effect of different sequencing depths across cells. Additionally, we also specified the following parameters: min.pct=0.1, min.diff.pct=0.1, logfc.threshold = 0, and only.pos = F. Bonferroni correction was applied for P-value adjustment using all peaks from each comparison, and peaks were considered significant if they had adjusted P-value<0.05.

#### Calculation of TF motif scores using snATAC-seq data

In order to evaluate TF binding accessibility profiles, we used the ChromVar tool^105^, which calculates biased-corrected deviations (TF motif scores) corresponding to gain or loss of accessibility for each TF motif relative to the average cell profile. To identify TFs with differential activity between cell groups of snATAC-seq data, we used a two-sided wilcoxon rank-sum test for the whole set of TFs in the assay and applied FDR correction for the resulting P-values. For subtype specific TFs we additionally required them to have positive Fold change in RNA-seq data when compared between tumor cells from each subtype vs pooled tumor cells from all other subtypes.

#### Identifying lineage-specific TF motifs in accessible chromatin regions using snATAC-seq data

To identify cancer lineage-specific motifs profiles in open chromatin, we first performed a comparison between tumor cells and their putative benign cell of origin (*i.e.*, basal-like BC cells and luminal progenitor cells, or luminal A/B cells and luminal mature cells) and all other epithelial cell groups (list of cell groups: basal-like BC, HER2-enriched BC, luminal A/B BC, luminal mature cells, luminal progenitor cells, and basal myoepithelial cells) using a two-sided Wilcoxon rank-sum test followed by FDR correction, as above. Cancer-associated non-lineage specific TFs were filtered out from this motif list by excluding TFs enriched in both luminal BC vs. luminal mature cells and basal-like BC vs. luminal progenitor cells. Individual differential motif accessibility comparisons were done for each BC subtype and benign cell type vs. all other epithelial cells, and any TFs significantly enriched in a non-lineage cell type (with mean motif score difference of at least 0.5) were also excluded from the lineage-specific motif list.

#### Visualizing the coverage of snATAC-seq data

For snATAC-seq coverage plots, we used the CoveragePlot function from the Signac package. For tumor samples, we plotted coverage for tumor cells only, and for normal cell populations we plotted coverage for combined cells across all samples.

#### Monocle pseudo-time analysis using snATAC-seq data

Trajectory-based analyses of snATAC-seq data were performed using Monocle2^106^. In order to build case-level trajectories, we used subsetted data from the snATAC-seq merged object. For each case we subsetted 1K cells from the pool of its tumor cells and also 1K cells from each of the combined sets of normal cell populations, e.g. luminal mature, basal/myoepithelial, and luminal progenitor. To create a monocle cds object we used the function newCellDataSet on slot count and top 50K expressed peaks, and with parameters lowerDetectionLimit=0.5 and expressionFamily=negbinomial.size(). To estimate size factors for each cell and dispersion function for the peaks we used functions estimateSizeFactors and estimateDispersions. Dimensionality reduction was performed using reduceDimension function with DDRTree method, and max_component=10.

#### scVarScan Mutation Mapping

We applied an in-house tool called scVarScan which can identify reads supporting the reference allele and variant allele covering the variant site in each individual cell by tracing cell and molecular barcode information in a scRNA bam file. The tool is freely available at https://github.com/ding-lab/10Xmapping. For mapping, we used high-confidence somatic mutations from WES data.

#### CNV calling on bulk whole exome data

Somatic copy number variants (CNVs) were called using GATK v4.1.9.0114. The hg38 human reference genome (NCI GDC data portal) was binned into target intervals using the PreprocessIntervals function, with a bin-length of 1,000bp and the interval-merging-rule set to OVERLAPPING_ONLY. A panel of normals (PON) was generated using each normal sample by utilizing the GATK functions CollectReadCounts with the argument --interval-merging-rule OVERLAPPING_ONLY, followed by CreateReadCountPanelOfNormals with the argument -- minimum-interval-median-percentile 5.0.

For tumor samples, reads that overlapped the target interval were counted using the GATK function CollectReadCounts. Subsequently, the tumor read counts were standardized and de-noised using the GATK function DenoiseReadCounts, with the PON specified by --count-panel-of-normals. Allelic counts for tumor samples were generated for variants present in the af-only-gnomad.hg38.vcf file, following GATK best practices (variants further filtered to 0.2 > af > 0.01 and entries marked with ’PASS’), using the GATK function CollectAllelicCounts. Segments were modeled using the GATK function ModelSegments, utilizing the denoised copy ratio and tumor allelic counts as inputs. Copy ratios for the segments were then called on the segment regions using the GATK function CallCopyRatioSegments.

To map the copy number ratios from segments to genes and assign amplifications or deletions, the Bedtools intersection ^107^ was used. For genes overlapping multiple segments, a custom Python script was employed to call that gene as amplified, neutral, or deleted based on a weighted copy number ratio calculated from copy ratios of each segment overlapped, the lengths of the overlaps, and the z-score threshold used by the CallCopyRatioSegments function. If the resulting z-score cutoff fell within the range of the default z-score thresholds used by CallCopyRatioSegments (0.9, 1.1), the bounds of the default z-score threshold were utilized instead, replicating the logic of the CallCopyRatioSegments function. Similarly, to map copy number ratios from segments to chromosome arms, another script was used following the same approach to call whether the chromosome arm was amplified, neutral, or deleted.

#### scRNA CNV Detection

To detect large-scale chromosomal copy number variations using single-cell RNA-seq data, inferCNV (version 0.8.2) was used with default parameters recommended for 10X Genomics data. All cells that are not tumor were pooled together for the reference normal set. inferCNV was run at a sample level and only with post-QC filtered data.

#### Differential scRNA Expression Analyses

For cell-level and cluster-level differential expression, we used the ’FindMarkers’ or ‘FindAllMarkers’ Seurat function as appropriate, with a minimum pct. of 0.25 and looking only in the positive direction, as lack of expression is harder to interpret due to the sparsity of the data. The resulting DEGs were then filtered for adjusted p-value < 0.05 and sorted by fold change. All differential expression analyses were carried out using the “SCT” assay.

#### Cell Surface Annotation

To annotate a given biomarker, we annotated each DEG by their subcellular location. Three databases were used to curate the subcellular location information: 1) Gene Ontology Term 0005886; 2) Mass Spectrometric-Derived Cell Surface Protein Atlas (CSPA)^108^; 3) The Human Protein Atlas (HPA) subcellular location data based on HPA version 19.3 and Ensembl version 92.38.

#### Identification of tumor markers using snRNA-seq data

In order to identify tumor cell surface markers we used functions of the Seurat package. We compared gene expression between tumor cells and non-tumor cells across snRNA-seq samples of Basal and Luminal subtypes. The pipeline consists of the following steps: (1) compare expression levels across cell types strictly within samples to discern markers characteristic of tumor cells, identifying those that hold more generally across all samples, (2) selecting the ones that were annotated as cell surface proteins by Gene Ontology (Term: 0005886). Using this approach, we identified candidate surface markers overexpressed in tumor cells compared to all the other cell types in a majority of individual samples (step 1) in each subtype. A gene is labeled tumor cell-specific if both the following criteria are satisfied: 1) the average expression of the gene is higher in tumor cells compared with any other cell type, respectively, for at least one sample, and that all the differences are of statistical significance (log(Fold Change) >0; adjusted P-value<0.05); 2) the average expression of the gene is higher in tumor cells compared with non-tumor cells (as a combined population) for 90% of the samples and that such differences are found to be statistically significant in at least 75% of the samples. All P-values were adjusted by Bonferroni correction.

#### Receptor Ligand Interactions

The CellPhoneDB Python package ^53^ was used to find interactions between cell types in individual objects. Annotation and input counts files were constructed as previously described^53^. Interactions were identified for scRNA-seq. The statistical analysis method of the CellPhoneDB package was run with 1000 iterations. Ligand-receptor pairs from the “significant means” output file were used in the downstream analysis. Interactions were filtered by the number of cells belonging to an interacting cell type (>10) and by the percentage of interacting cells in the total number cells in a sample (> 0.1%). Only interactions annotated as “curated” were used for the analysis.

#### Differential gene expression analysis for subtype specific transcriptional changes

We utilized the Seurat function FindMarkers (test.use = ’LR’, only.pos = T, logfc.threshold = 0.2, min.pct = 0.1) to evaluate differentially expressed genes (DEGs) between epithelial cell types. First we identified genes that were differentially expressed between related lineages against other epithelial cell types including the following groups: (G1) basal-like tumor and luminal progenitor vs. all other epithelial cell types; (G2) luminal A tumor, luminal B tumor, luminal mature vs. all other epithelial cell types; (G3) luminal progenitor and luminal mature vs. all other epithelial cell types. Second we identified cancer specific genes that were upregulated in the specific subtype relative to their cell of origin by determining DEGs for the following groups: (C1) basal-like tumor vs. luminal progenitor; (C2) luminal A tumor and luminal B tumor vs. luminal mature. Finally we extracted all DEGs (A1) for each epithelial cell type vs. all other cells in the epithelial subset (FindAllMarkers). This analysis was performed only on the single-nuclei RNA-sequencing data. For analyzing the changes between basal tumors and luminal progenitors we focused our analysis on genes identified in G1 with p_val_adj < 0.05. We then filtered out genes that were found in C1 or C2, being only cancer type related changes and not related to cell of origin. We then filtered out genes in G2, DEGs related to the luminal tumor transition. Using the all DEG analysis (A1) we extracted DEGs specific to basal tumor or luminal progenitor and filtered out any genes specific to luminal A tumor, luminal B tumor, HER2 tumor, basal/myoepithelial and luminal mature DEG analysis. Using this final gene list we extracted the average expression across all epithelial cell types (focusing on genes with pct.exp>20 and avg_log2FC > 1) for our final analysis. For analyzing the changes between luminal A/B tumors and luminal mature cells we focused our analysis on genes identified in G1 with p_val_adj < 0.05. We then filtered out genes that were found in C1 or C2, being only cancer type related changes and not related to cell of origin. We then filtered out genes in G1, DEGs related to the basal-like tumor transition. Using the all-DEG analysis (A1) we extracted DEGs specific to luminal A tumor, luminal B tumor, or luminal mature and filtered out any genes specific to HER2 tumor, basal/myoepithelial or basal-like tumor DEG analysis. Using this final gene list we extracted the average expression across all epithelial cell types (focusing on genes with pct.exp>20 and avg_log2FC > 1) for our final analysis.

#### Analysis of Lymphocyte Dense Clusters in Spatial Transcriptomics Data

For each spatial transcriptomics spot overlapping a lymphocyte dense region (Extended Data Fig. 2), spots were subset from the RDS object for each sample and merged using the Seurat merge function. At least three lymphocyte dense regions were identified on each slide.

#### Spatial mapping of snRNA-seq cell types to Spatial Transcriptomics Data

We generated a joint cell type reference Seurat object for each PAM50 subtype by merging snRNA-seq data from the same subtype (Basal, LumA, LumB, and Her2). Using these snRNA-seq references, we then infer the cell type composition in Spatial Transcriptomics (ST) samples of the same PAM50 subtype using CytoSPACE ^54^, a tool that align snRNA-seq to ST data and resolve cell type compositions per spatial spot. A custom script was developed to facilitate preprocessing the snRNA-seq and ST file, as well as integrating the CytoSPACE result into the Seurat object for easier downstream analysis and visualization. These workflow and processing scripts can be found in the Github page associated with this manuscript.

### CODEX Quantification and Analysis

#### Multiplex image segmentation

Multiplex images were segmented using the Mesmer pretrained nuclei + membrane segmentation model in the Deepcell ^109^ cell segmentation library. The DAPI channel was used as the nuclei segmentation image, and Pan-Cytokeratin, E-cadherin, CD45, CD8, CD3, Vimentin, SMA, CD31, and CD20 channels were merged and used as the membrane segmentation image. Following segmentation, cells were classified as positive or negative for the following epithelial markers: GATA3, cKit, CK14, CK19, ER, PR, and Her2. To eliminate batch effects, marker thresholds were set manually for each image.

#### Normal/tumor region identification and classification

We then identified epithelial regions in an unsupervised fashion. First, Pan-Cytokeratin and E-cadherin were thresholded on the values discussed in the section above. The masks were then merged into a consensus mask. This mask was passed through a gaussian filter (sigma=2.0) and hole-filling algorithm. The regions in the resulting mask (n=12,513 across all images) were then classified into normal and tumor regions.

A pseudo-color RGB image was generated for each region that represented image intensities for Pan-Cytokeratin, SMA, DAPI, and Podoplanin. A subset of these regions (n=637) were then manually annotated as normal, ductal carcinoma in-situ, invasive ductal carcinoma, or image artifacts. These annotations were then partitioned via a 80/20 split into training and validation datasets. During model training, images were augmented by random affine rotation and color jitter. A convolutional neural network was used to classify these images based on region type. The neural network consisted of two ResNet34 stems (one for the 3-channel pseudo-color RGB image, and another for a 1-channel mask representing the pixels the bounds of the region in the pseudo-color RGB image), the stems are then merged into a prediction head consisting of three linear layers separated by ReLU activation functions and batch normalization layers. The final layer is followed by a Softmax activation that outputs classification probabilities. The network was trained for 500 epochs and achieved a validation accuracy of 96%. The network was then used to predict all 12,513 regions. All regions predicted by the model were then manually reviewed to further reduce the region annotation error rate. In total, 649 regions were annotated as normal, and 10,753 as tumor. Images classified as image artifacts were excluded from downstream analysis.

#### Epithelial marker comparison among normal and tumor regions across subtypes

For epithelial marker comparisons, the fraction of positive cells within each tumor region for each epithelial marker was calculated. Values for each region were then averaged at the sample level prior to plotting and significance calculations.

### Immunofluorescence Quantification and Analysis

Immunofluorescence images were first standardized by considering only the first 1250 lines of each image. In order to quantify MELK signal, a mask based on e-cadherin was generated by adaptive thresholding. For each sample, the threshold value was set to half of the e-cadherin sample mean. The mask was then generated as follows: for a given sample, it was set to 1 where the e-cadherin value was above the threshold, and set to 0 elsewhere. The average of MELK pixel intensity per sample was then calculated considering only the positive regions of the mask. Finally, the resulting averages were grouped by their case (5 each) and the results were displayed in a violin plot. Corresponding p-values between all basal-like and luminal cases were calculated.

## Data and Code Availability

All human sequencing and imaging data are being deposited via the Human Tumor Atlas Network (HTAN) dbGaP Study Accession: phs002371.v1.p1 (https://www.ncbi.nlm.nih.gov/projects/gap/cgi-bin/study.cgi?study_id=phs002371.v1.p1). In addition, all data will be deposited to the HTAN Data Coordinating Center Data Portal at the National Cancer Institute: https://data.humantumoratlas.org/ (under the HTAN WUSTL Atlas). Scripts related to specific analysis in the paper can be found at the following github link: https://github.com/ding-lab/HTAN_BRCA_publication/tree/main. All mouse snRNA/snATAC data has been deposited to Gene Expression Omnibus under series GSE240577.

