## Extended Data Figures for "Differential chromatin accessibility and transcriptional dynamics define breast cancer subtypes and their lineages"

### Extended Data Fig. 1

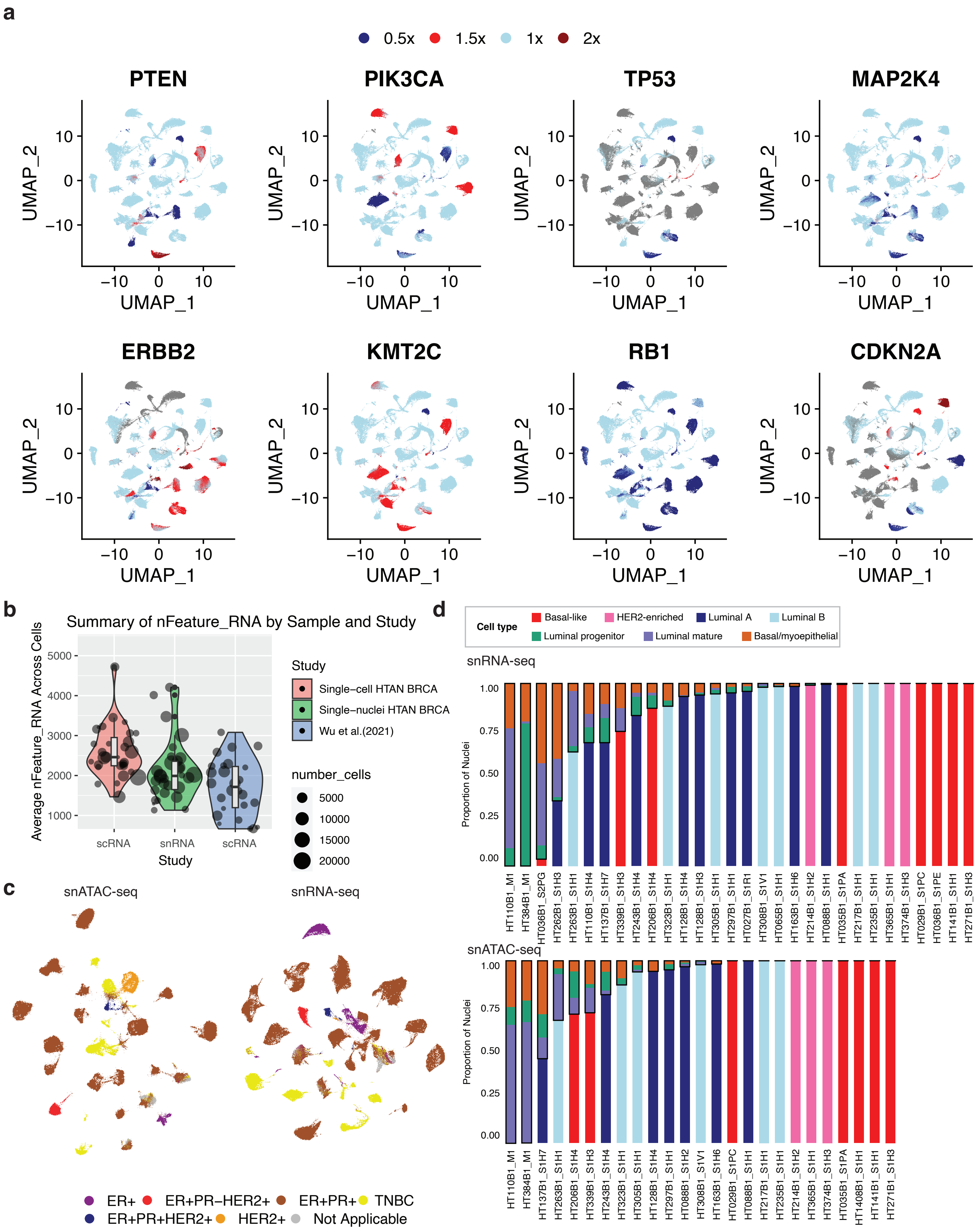

### Extended Data Fig. 2

**a**

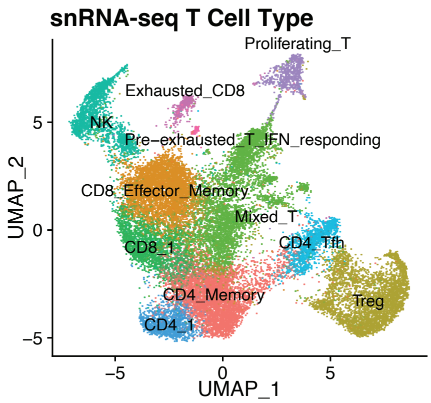


Wilcoxon Test Result

| T Cell Type | Group 1 | Group 2 | P |
|---|---|---|---|
| CD4_Memory | Luminal | Basal | 0.04609563 |
| CD4_Tfh | Luminal | Basal | 0.04609563 |
| Exhausted_CD8 | Luminal | Basal | 0.03149074 |

**b** **c**

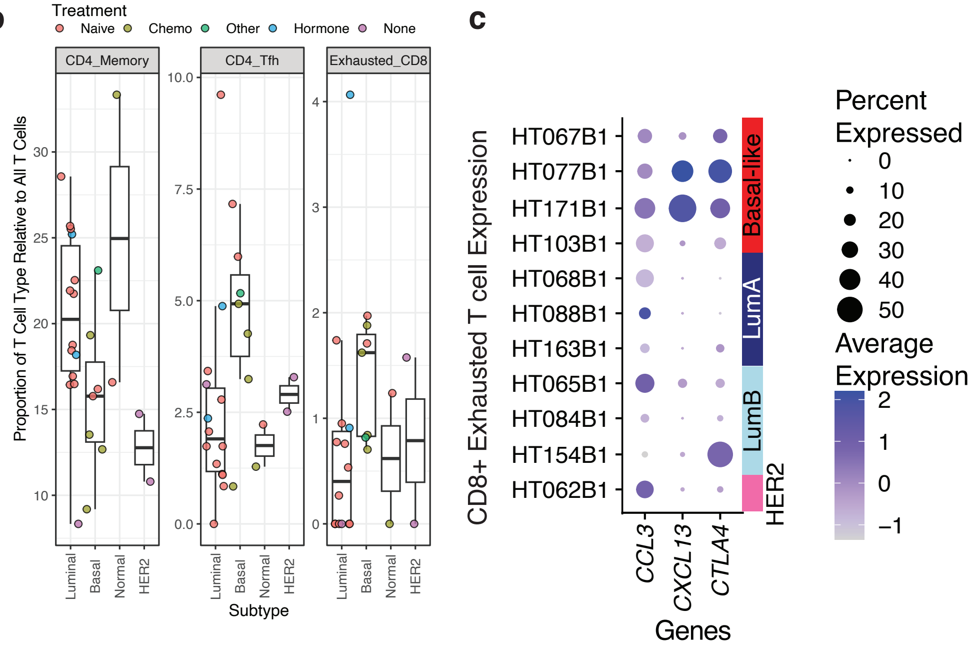

### Extended Data Fig. 3

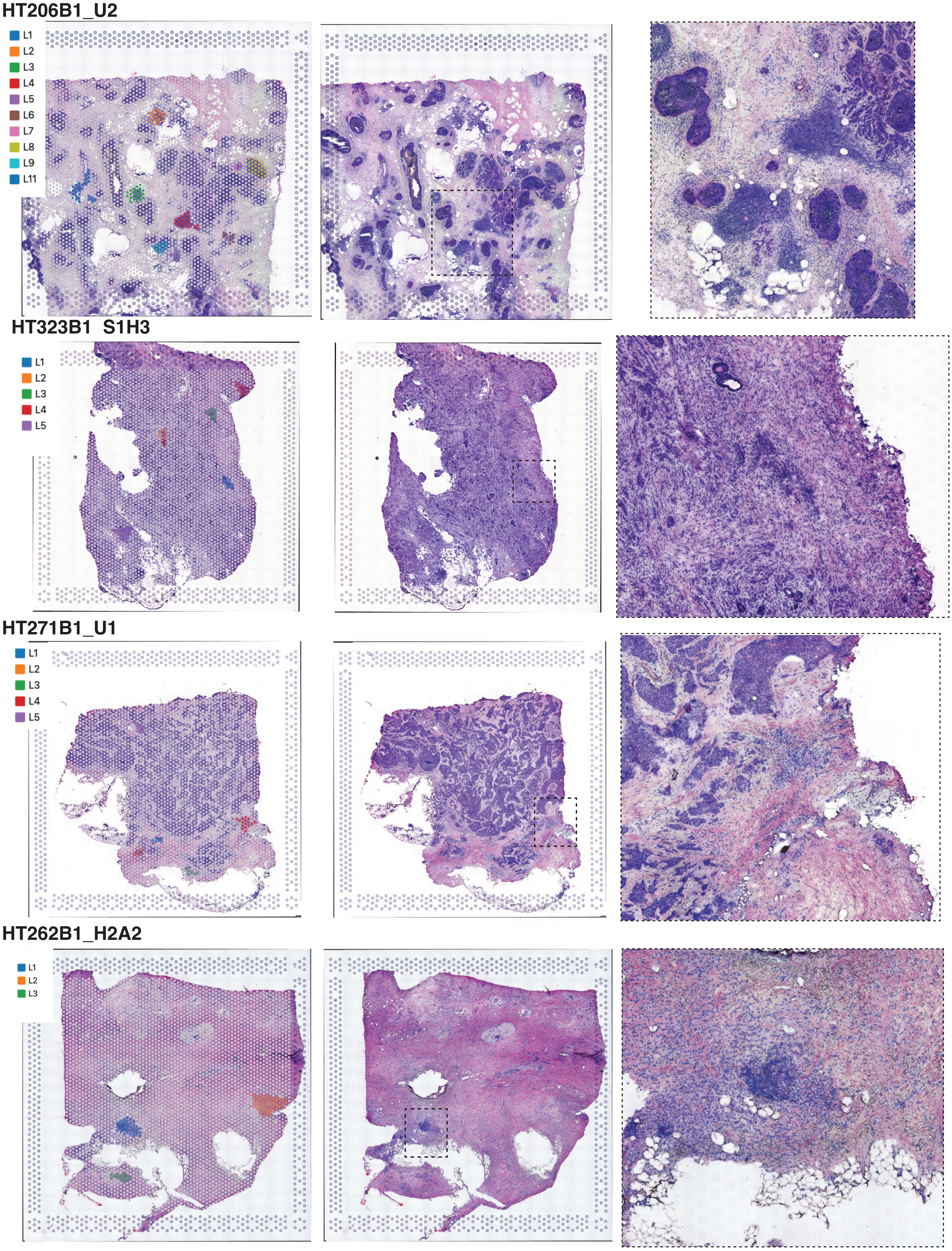

### Extended Data Fig. 4

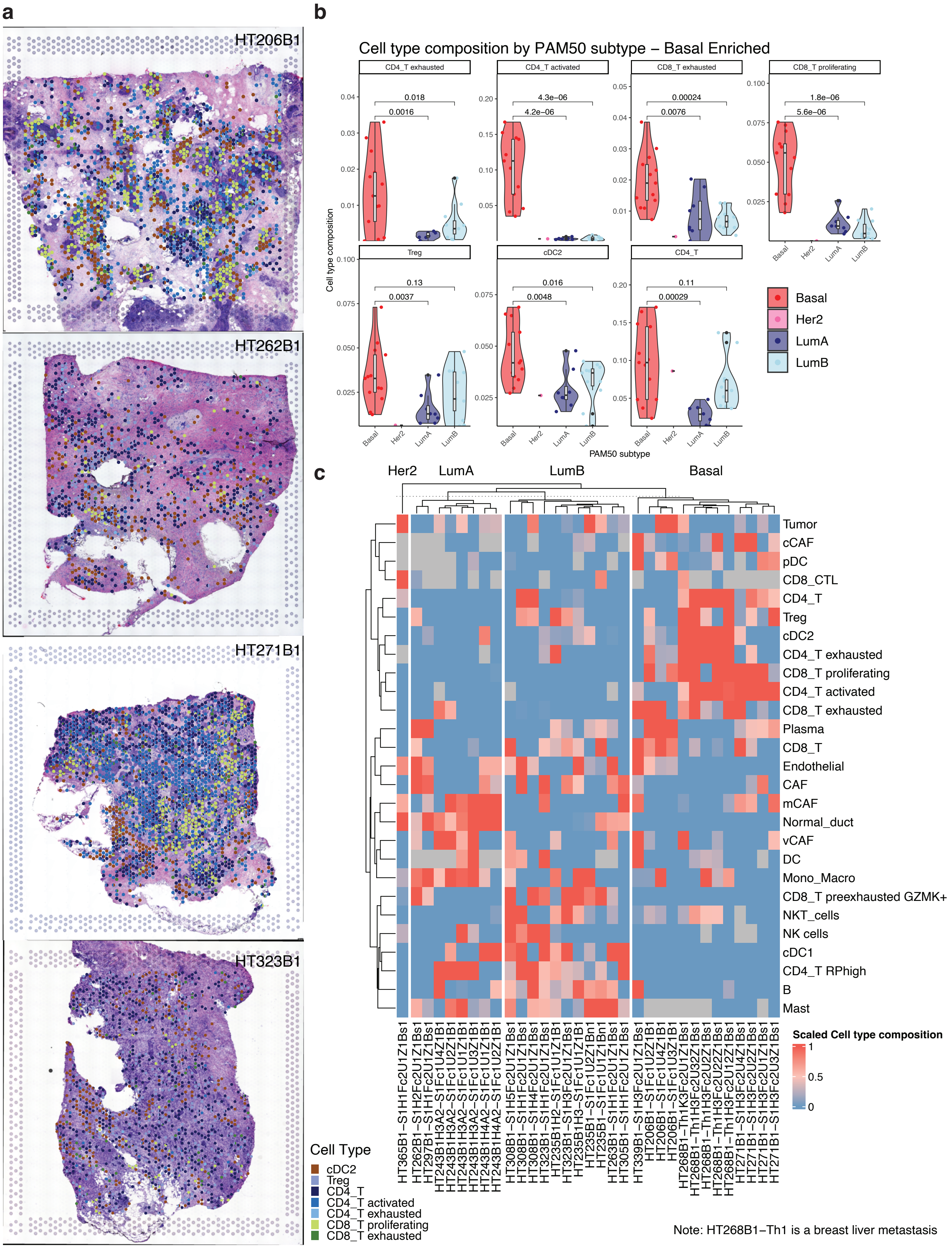

### Extended Data Fig. 5

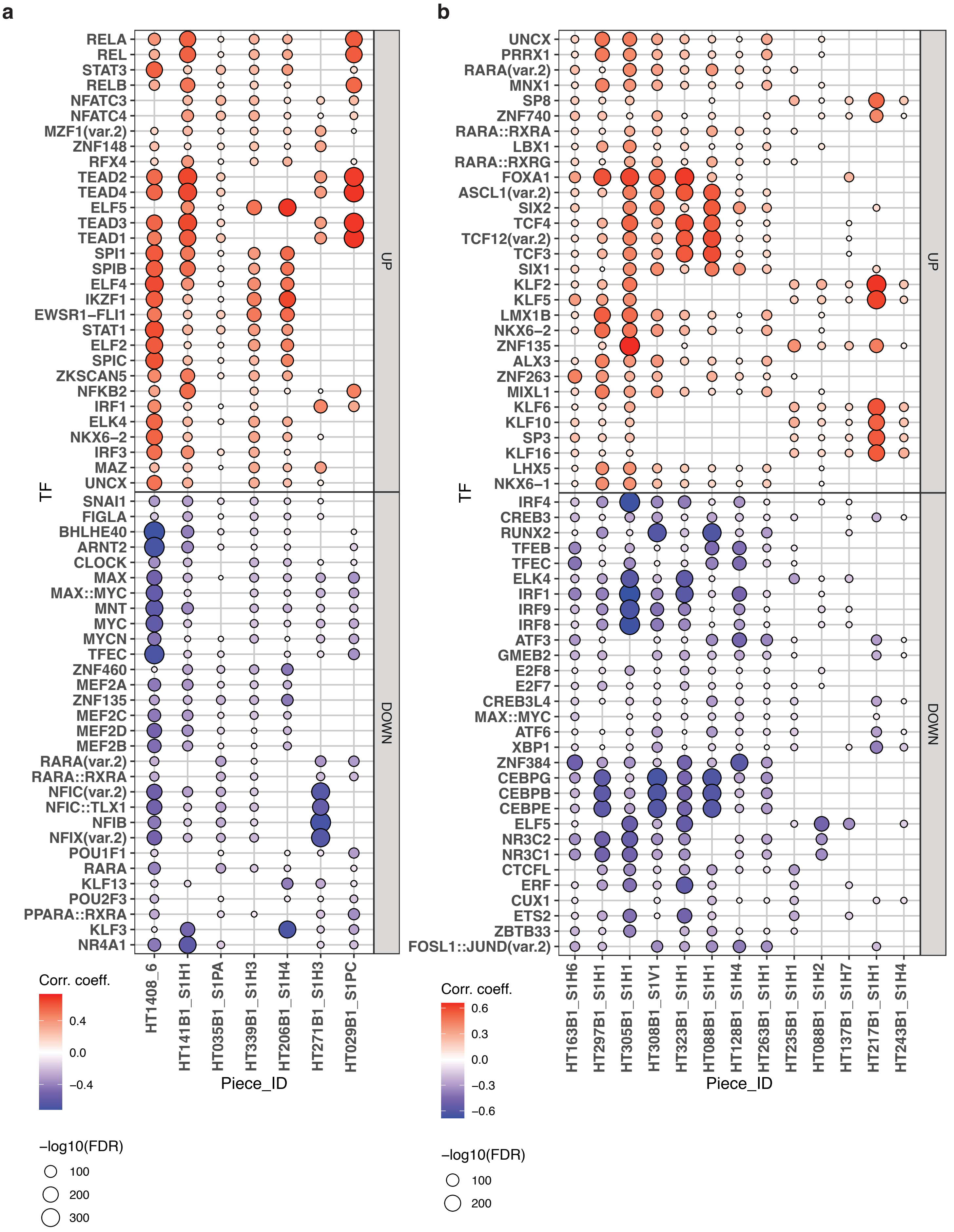

### Extended Data Fig. 6

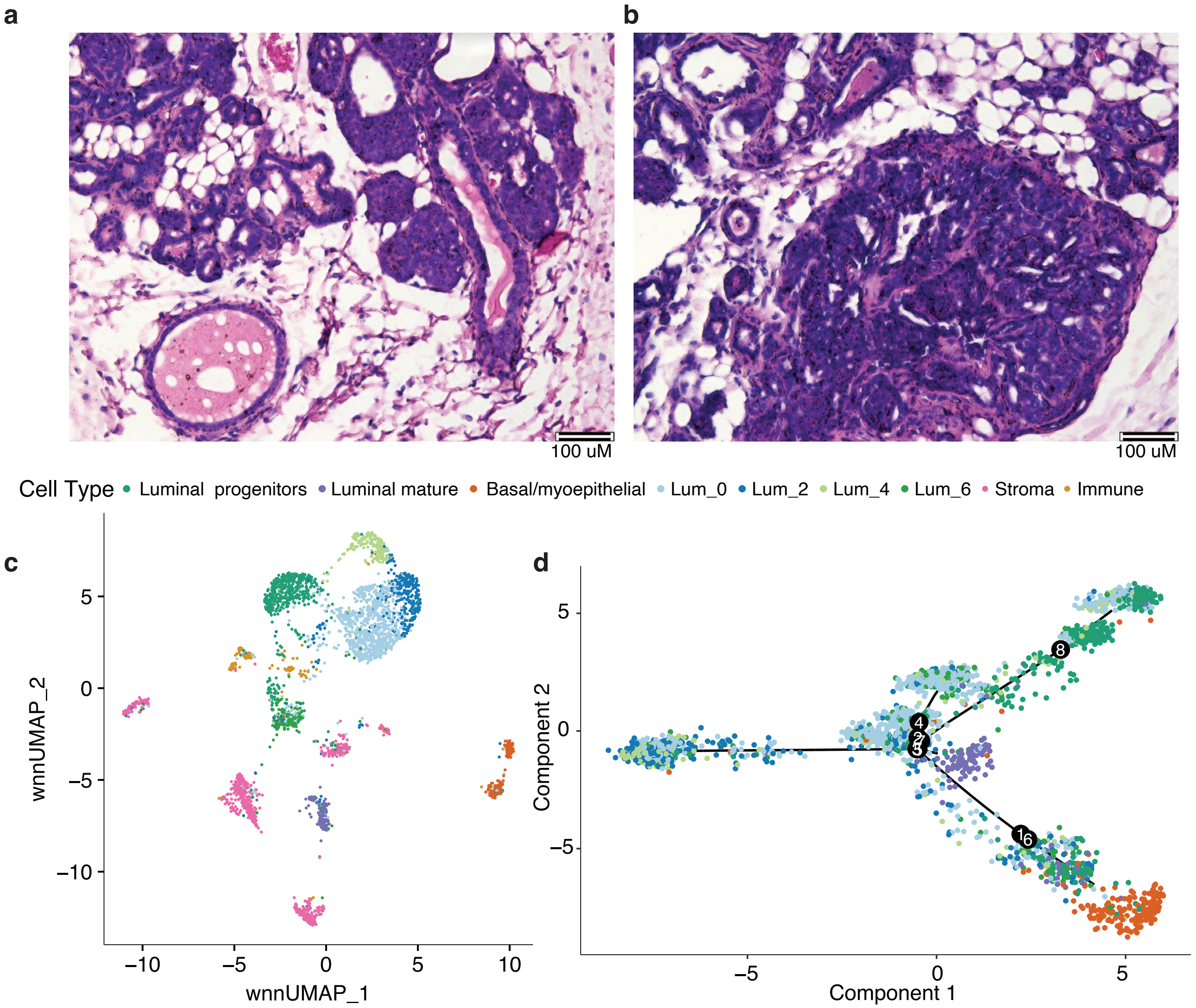

### Extended Data Fig. 7

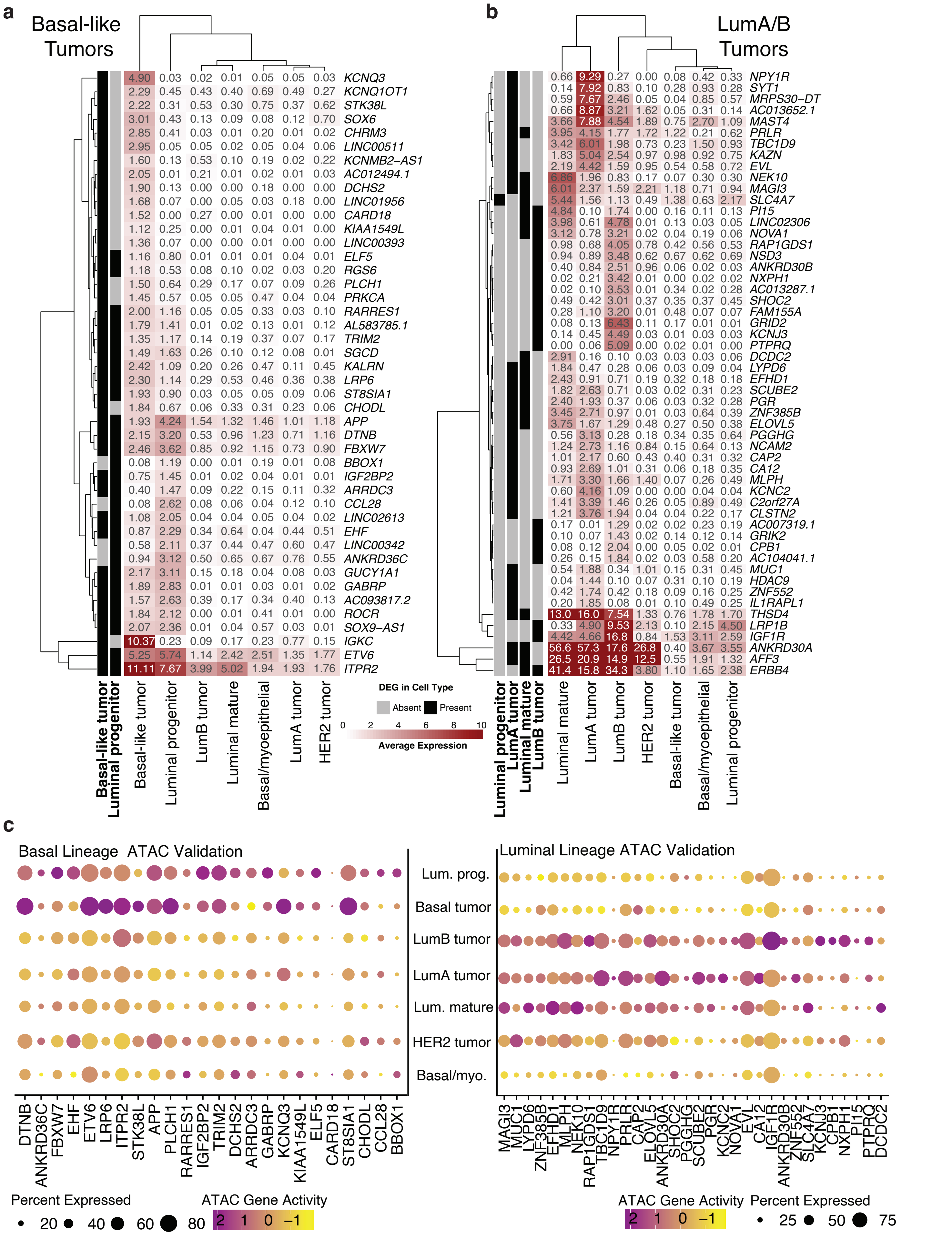

### Extended Data Fig. 8

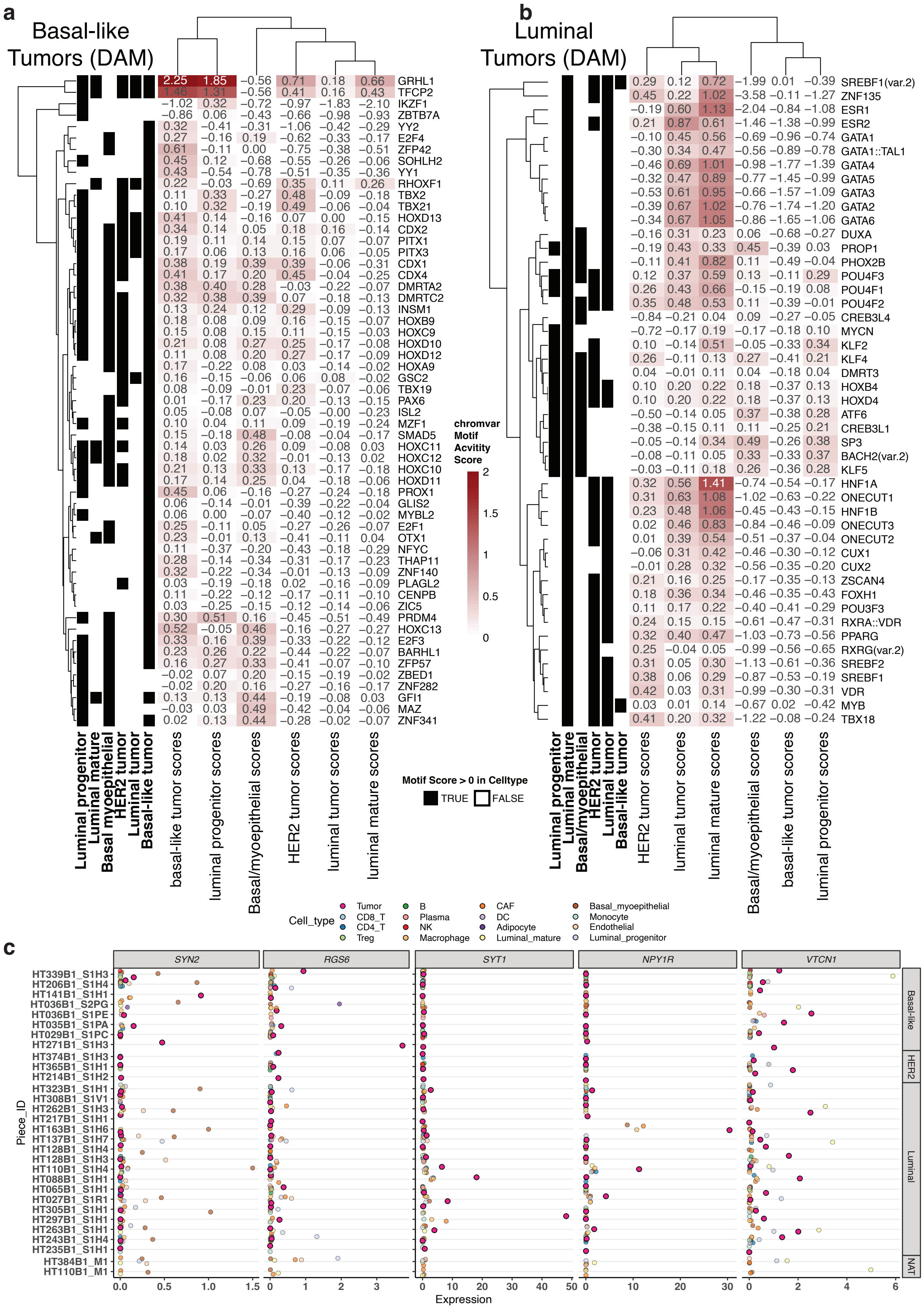

### Extended Data Fig. 9

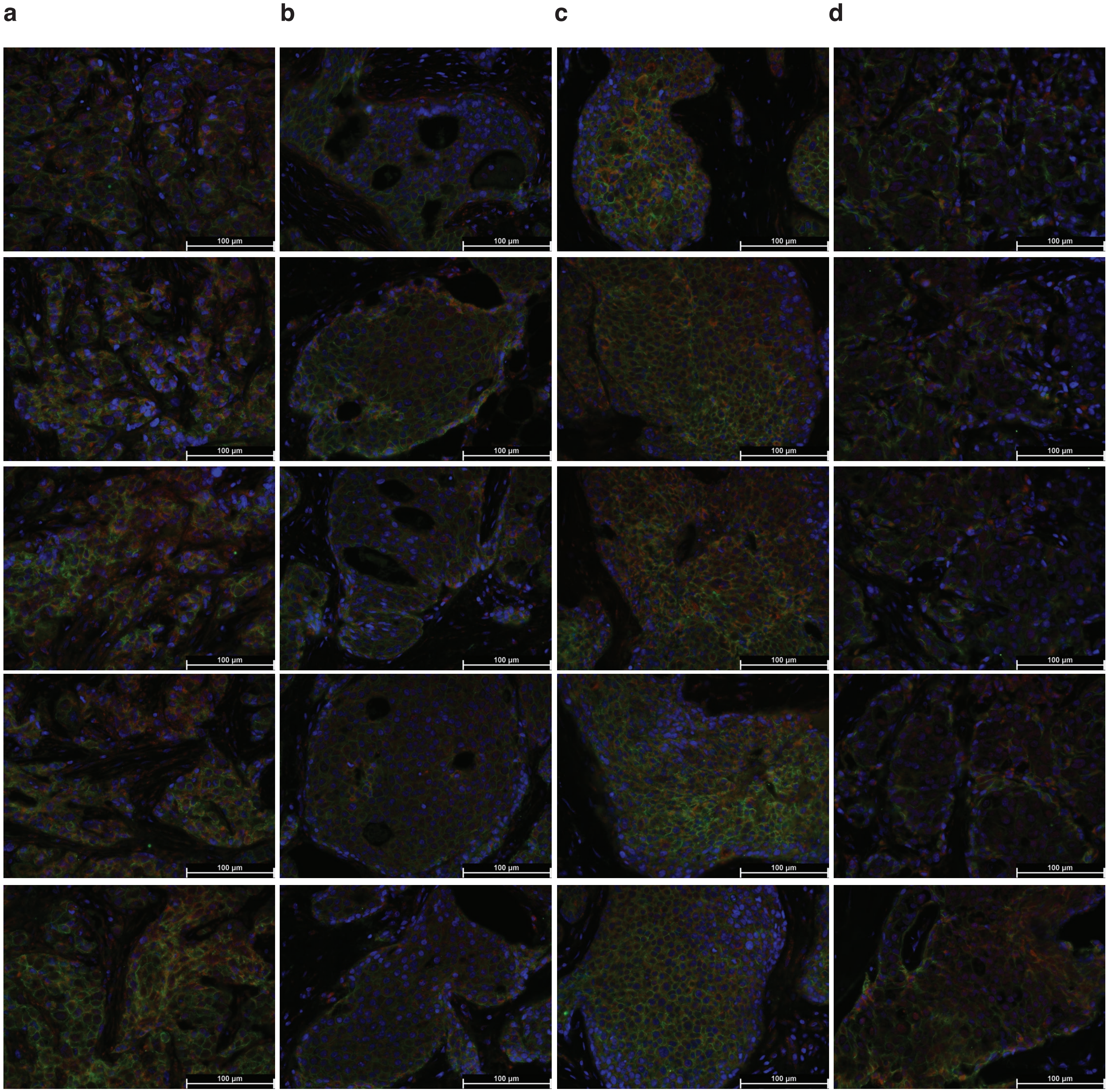

### Extended Data Fig. 10

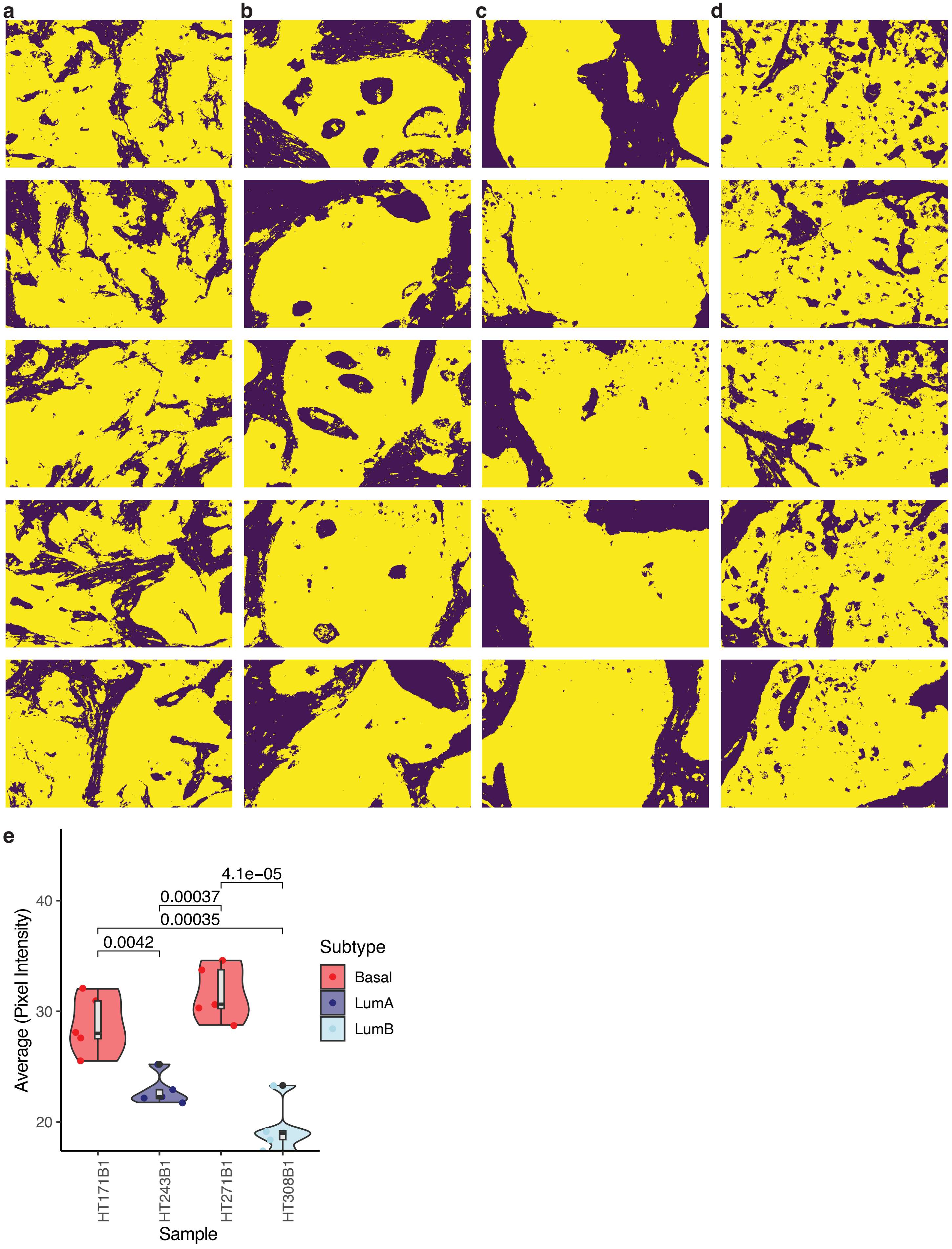
